# H3AGWAS : A portable workflow for Genome Wide Association Studies

**DOI:** 10.1101/2022.05.02.490206

**Authors:** Jean-Tristan Brandenburg, Lindsay Clark, Gerrit Botha, Sumir Panji, Shakuntala Baichoo, Christopher Fields, Scott Hazelhurst

## Abstract

**Background:** Genome-wide association studies (GWAS) are a powerful method to detect associations between variants and phenotypes. A GWAS requires several complex computations with large data sets, and many steps may need to be repeated with varying parameters. Manual running of these analyses can be tedious, error-prone and hard to reproduce.

**Results:** The H3AGWAS workflow from the Pan-African Bioinformatics Network for H3Africa is a powerful, scalable and portable workflow implementing pre-association analysis, implementation of various association testing methods and post-association analysis of results.

**Conclusions:** The workflow is scalable — laptop to cluster to cloud (e.g., SLURM, AWS Batch, Azure). All required software is containerised and can run under Docker on Singularity.

## Background

Genome-wide association studies (GWAS) are a powerful method to detect associations between variants and phenotypes; from initial raw genotype data until detection of putative causal variant requires numerous steps, software and approaches to extract and understand results [1]. Common steps after genotyping include:

1. Preparing data into standard formats

2. Quality control (QC) of genotypes to remove uncertain positions and individuals – e.g., discrepancy between genotyped sex and known sex, and bias due to high relatedness between individuals. These are important steps to reduce noise and false positive discovery rate [2–4].

3. Associating genetic variation with phenotype. This step is very expensive, with millions of positions and sample sizes ranging from several thousands to several hundred thousand. These methods take account of relatedness between individuals with mixed models and different algorithms to improve detection and/or approximations for a very large sample size.

4. Post-association analysis, which may include highly complex methods such as meta analysis considering different GWAS summary statistics, fine-mapping to define causal variants, heritability of phenotype, replication and transferabilty of previous results, annotation, integration of eQTL, and/or calculation of polygenic risk score [5].

## Motivation

The phases of GWAS are all complex, and typically require multiple executions, sometimes on different platforms by different collaborators and replicability of analyses is crucial. The PanAfrican Bioinformatics Network of the Human Heredity and Health in Africa Consortium [6] (H3ABioNet) has as one of its goals the task of supporting the work of H3Africa, and African scientists more broadly by developing workflow for commonly performed analyses. Baichoo et al. [7] provide an overview of workflow development within H3ABioNet, including an introduction to a much earlier version of this workflow. The goal of the H3AGWAS workflow is to support scientists undertaking GWAS taking into account access to heterogeneous computing environments.

In summary, the goal of the H3AGWAS workflow is to provide a flexible, powerful and portable workflow for genome-wide association studies. The use of a workflow reduces the manual intervention required by human analysts, thereby reducing the overall time for a project to complete. Some phases of a GWAS are exploratory and analyses may need to be re-run as QC proceeds, and different parameters and analytic techniques tried after assessing initial results. The workflow needs to support reproducible analyses and be portable and scalable across many different computational environments (laptop to cluster to cloud), reflecting the heterogeneous environments across Africa. Using Nextflow and containerisation promotes scalability and portability.

## Implementation

The workflow has been developed in Nextflow [8], with Python [9], bash and R scripts [10] and uses well-known bioinformatics tools. It can easily be ported to different execution environments (e.g., standalone, job scheduling, cloud) and uses containers to package software and dependencies assures replicability and simple installation. Figure 1 gives an overview.

**Figure 1.**
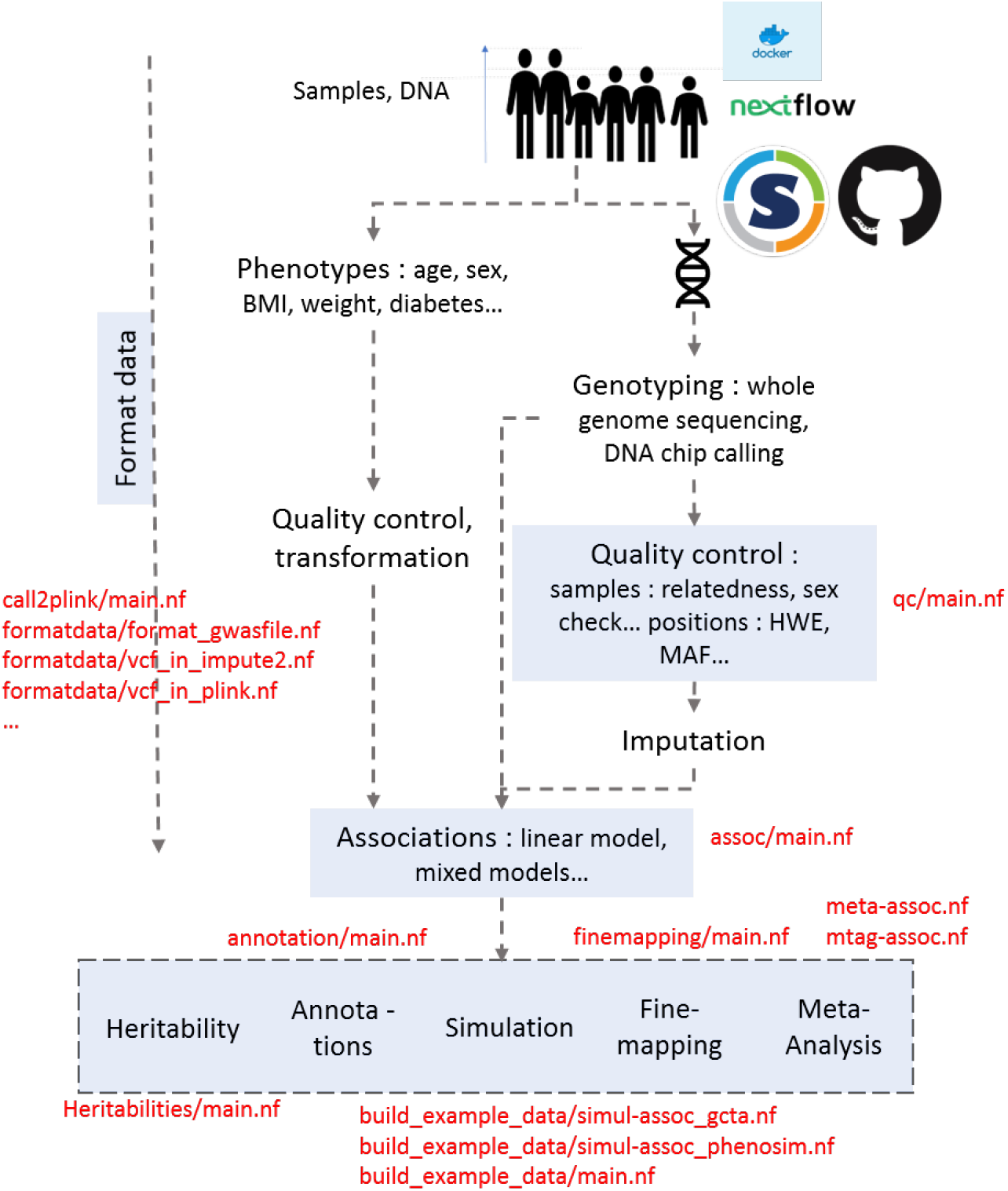
Overview of GWA studies from DNA sample until post-association analysis, box in blue corresponding to part of GWA studies present in H3AGWAS workflow and text in red corresponding to scripts in H3AGWAS workflow

Rather than producing one workflow which operates end-to-end, the H3AGWAS workflow is split into several independent sub-workflows mapping to separate phases of work. Independent workflows allow users to execute parts that are only relevant to them at those different phases. For example, our experience has shown that the QC step requires multiple iterations over several weeks to find the best QC parameters and resolve problems with data. Once the QC is complete, the analysis moves to the next phase, which in turn may take weeks. Sample runs and extensive documentation for the different phases can be found at http://github.com/h3abionet/h3agwas/.

### Pre-association Workflows

#### Producing PLINK data

The `call2plink` workflow converts Illumina genotyping reports into PLINK format.

#### Quality control

The `qc` workflow performs quality control on a set of input PLINK file. The workflow considers per-sample and per-single nucleotide polymorphism (SNP) missingness, minor allele frequency, levels of heterozygosity, highly related samples, possible duplicates, and sex mismatches, and also examines possible batch effects (for example, between cases and controls, for samples collected from different sites, or genotyped in different runs). A detailed report is produced which helps the user understand the data and which can be used in the methods section of a paper. All QC and workflow parameters (including software versions) and the MD5 checksums of input and output data are recorded in order to promote replicability and reduce the risk of version skew.

#### Association testing

The `assoc` workflow performs association on PLINK formatted files, including adjustment for multiple testing in PLINK. In addition to the basic association tests, the workflow currently supports Cochran-Mantel-Haenszel (CMH), linear and logistic regression, permutation and mixed-model association testing. This workflow provides user-selectable choices of software for association testing. PLINK is the work-horse for basic linear models, including support of covariates and adjusting for population structure. Exact linear mixed models with relatedness matrix have been included (Fast-LMM [11] and GEMMA [12]). For larger data sets, BOLTLMM [13] and fastGWA [14, 15], SAIGE [16] and regenie [17] which use approximation of relatedness can be selected (and the workflow can compute the SNP-derived genetic relationship matrix (GRM) from genotype data using GCTA [15]). Besides PLINK format, BOLT-LMM, SAIGE, fastGWA and regenie also accept dosage as optional input (e.g., for imputed data). BGEN format can be extracted from VCF files after imputation, using formatting scripts (see the *Format conversion* section below) – the `assoc` pipeline supports these formats.

Many common complex traits are believed to be a result of the combined effect of genes, environmental factors and their interactions. Gene–environment interaction (G *×* E) can be analysed to detect loci where genotype-phenotype association may depend on the environment: G *×* E options from GEMMA and PLINK are implemented in the workflow (see Figure 2 and Table 1).

**Figure 2.**
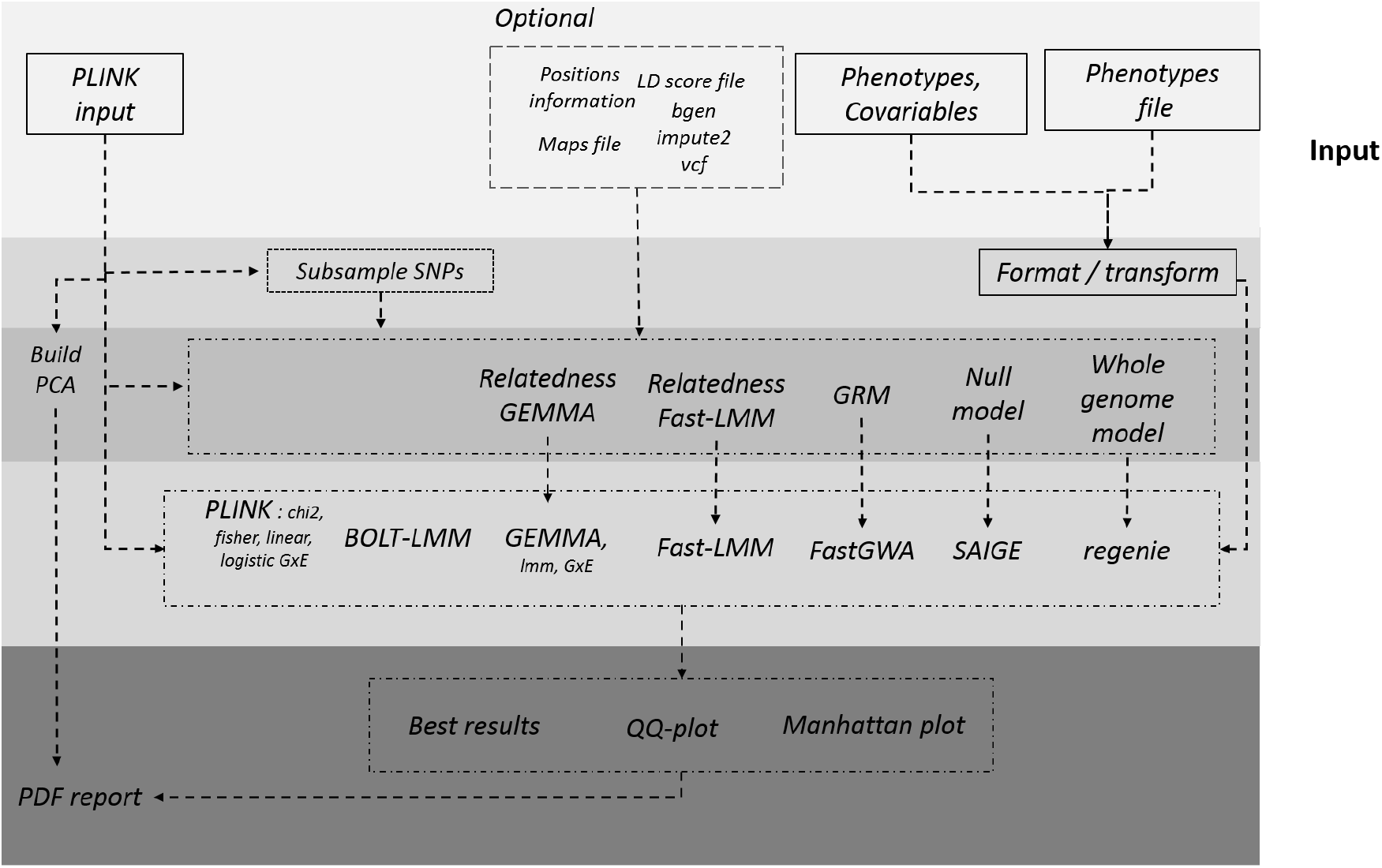
Workflow of association testing each background color represent different steps from input to output with preparation of input data, generated relatedness matrix or GRM to take account population structure and association testing

**Table 1:** List of softwares and resources used in H3AGWAS workflow, softwares are classify by phase of GWAS and task.

| Phase | Software/Resource | Workflow / Task | Ref |
| --- | --- | --- | --- |
| All | PLINK (1.9) | All | [18] |
|  | Python (3) | All | [9] |
|  | R (3.6) | Plot and extraction data | [10] |
| Association testing | GEMMA (0.98.5) | Association testing / heritability / conditional analysis | [12] |
|  | Fast-LMM (binary version) | Association testing | [11] |
|  | BOLT-LMM | Association testing / heritability | [13] |
|  | SAIGE 1.0 | Association testing | [16] |
|  | regenie 3.1.3 | Association testing | [17] |
|  | GCTA : fastGWA | Association testing | [14, 15] |
| Post-association analysis | GCTA : COJO-slct, simulation, GREML | Fine-mapping / heritability/ simulation | [19] |
|  | MetaSoft | Meta-analyses | [20] |
|  | GWAMA | Meta-analyses | [21] |
|  | METAL | Meta-analyses | [22] |
|  | MTAG | Multi-trait association | [23] |
|  | LDSC | Heritability | [24] |
|  | PhenoSim | Simulation | [25] |
|  | LocusZoom | Annotations | [26] |
|  | Annovar | Annotations | [27] |
| Format data | BCFtools | Convert vcf | [28] |
|  | QCTOOL (v2) | Convert vcf to bgen/bimbam/impute2 | [29] |
|  | CrossMap | Convert positions between genome builds | [30] |

The PLINK input files are also used to perform a principal component analysis (PCA) and a PCA plot is generated that can be used to identify any possible population structure in the data set.

Output includes a report with PCA, Manhantan plot, qq plot of each phenotype, summary statistics and software versions used by the pipeline.

#### Post-association analysis

The post-association analysis workflows use genotype data and results of association testing in order to (1) find putative causal variants; (2) perform a meta-analysis or multi-trait genomewide association study using summary statistics; (3) estimate global heritability; and (4) annotate positions (see Tables 1 and 2).

**Table 2:**
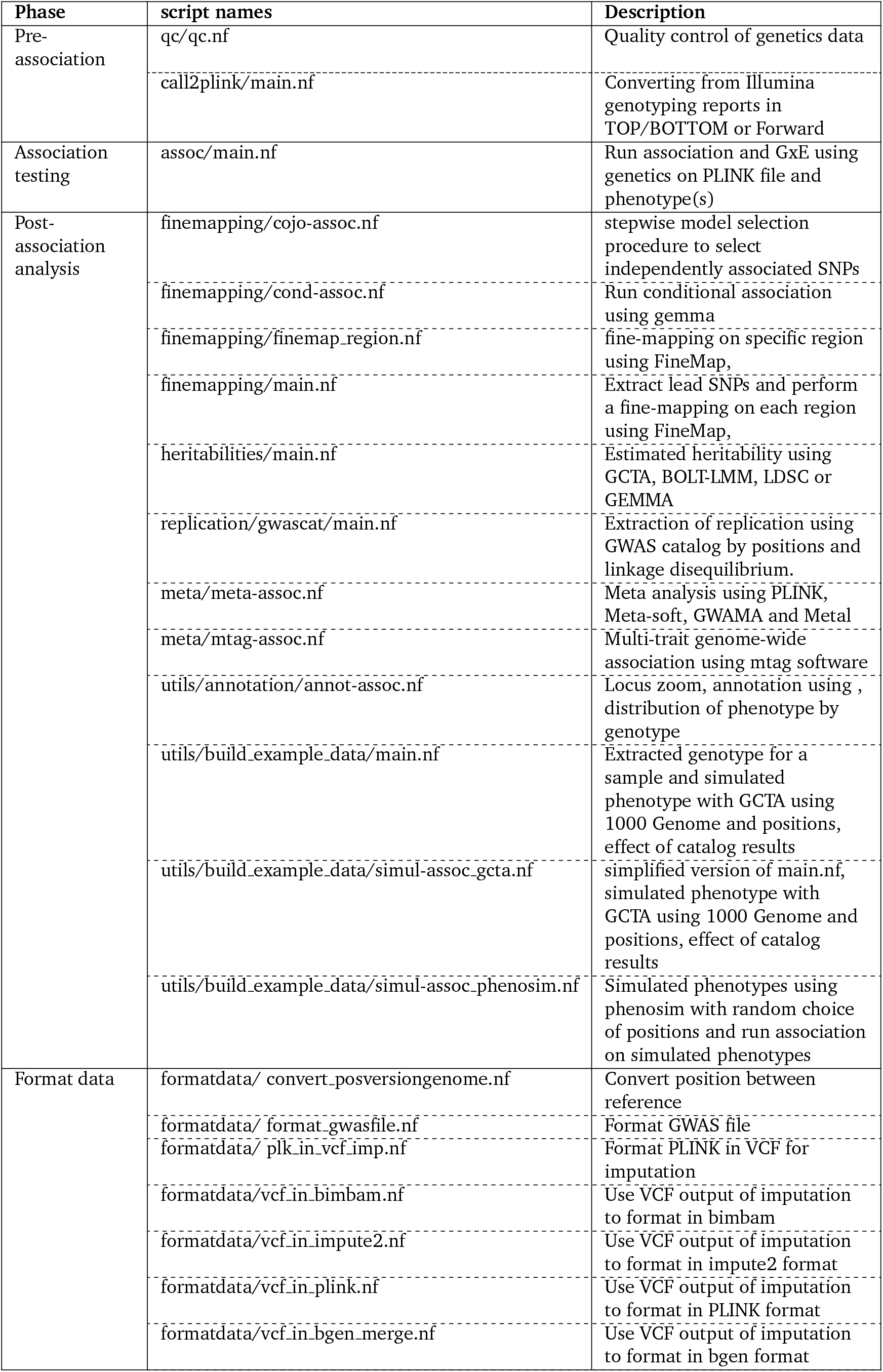
List and description of Nextflow scripts by phase of GWAS

| Phase | script names | Description |
| --- | --- | --- |
| Pre-association | qc/qc.nf | Quality control of genetics data |
|  | call2plink/main.nf | Converting from Illumina genotyping reports in TOP/BOTTOM or Forward |
| Association testing | assoc/main.nf | Run association and GxE using genetics on PLINK file and phenotype(s) |
| Post-association analysis | finemapping/cojo-assoc.nf | stepwise model selection procedure to select independently associated SNPs |
|  | finemapping/cond-assoc.nf | Run conditional association using gemma |
|  | finemapping/finemap_region.nf | fine-mapping on specific region using FineMap, |
|  | finemapping/main.nf | Extract lead SNPs and perform a fine-mapping on each region using FineMap, |
|  | heritabilities/main.nf | Estimated heritability using GCTA, BOLT-LMM, LDSC or GEMMA |
|  | replication/gwascats/main.nf | Extraction of replication using GWAS catalog by positions and linkage disequilibrium. |
|  | meta/meta-assoc.nf | Meta analysis using PLINK, Meta-soft, GWAMA and Metal |
|  | meta/mtag-assoc.nf | Multi-trait genome-wide association using mtag software |
|  | utils/annotation/annot-assoc.nf | Locus zoom, annotation using , distribution of phenotype by genotype |
|  | utils/build_example_data/main.nf | Extracted genotype for a sample and simulated phenotype with GCTA using 1000 Genome and positions, effect of catalog results |
|  | utils/build_example_data/simul-assoc_gcta.nf | simplified version of main.nf, simulated phenotype with GCTA using 1000 Genome and positions, effect of catalog results |
|  | utils/build_example_data/simul-assoc_phenosim.nf | Simulated phenotypes using phenosim with random choice of positions and run association on simulated phenotypes |
| Format data | formatdata/ convert_posversiongenome.nf | Convert position between reference |
|  | formatdata/ format_gwasfile.nf | Format GWAS file |
|  | formatdata/ plink_in_vcf_imp.nf | Format PLINK in VCF for imputation |
|  | formatdata/vcf_in_bimbam.nf | Use VCF output of imputation to format in bimbam |
|  | formatdata/vcf_in_impute2.nf | Use VCF output of imputation to format in impute2 format |
|  | formatdata/vcf_in_plink.nf | Use VCF output of imputation to format in PLINK format |
|  | formatdata/vcf_in_bgen_merge.nf | Use VCF output of imputation to format in bgen format |

#### Genetic heritability and co-heritability of phenotypes

There are two scripts to compare heritability and co-heritability of phenotypes. The first uses relatedness and phenotypes, based on REML or variance components analysis with BOLT-LMM [13, 31], GEMMA [12] and GCTA [14, 32]. The second uses summary statistics and methods implemented in GEMMA [33] and LDSC [24]. Furthermore, the workflow can compute the co-variability and coheritability between phenotypes using relatedness and phenotypes using BOLT-LMM and GCTA or with summary statistics with LDSC.

#### Annotations

An annotation script extracts genotypes of each individual and compares to phenotype, annotates lead SNPs using Annovar [27], plots regions using LocusZoom software [26], plot distribution of phenotypes by genotypes, and generates a report for the user.

#### Simulations of phenotypes

To estimate true and false positive detection in GWA, `simul-assoc_ phenosim.nf` script randomly builds phenotypes using the PhenoSim software [25] and genetics data where loci are randomly selected, followed by a GWA on the simulated data using BOLT-LMM and GEMMA (see supplementary table 7). In addition, the build_example_data/main.nf script builds phenotypes of individuals using initial genotype and allele effects. By default, the workflow uses 1000 Genomes Project (KGP) data [34] and extracts effect of positions from the GWAS Catalog. The steps are: (1) extract and format KGP data; (2) download GWAS catalog positions and results; (3) simulate phenotype in KGP individuals using effect of position, using GCTA [18] (see supplementary table 7).

#### Causal variants

Three workflows have been implemented to detect causal variants: finemapping/cojo-assoc.nf uses a step-wise model selection procedure to select independently associated SNPs [19]. `finemapping/main.nf` or `finemapping/finemap_region.nf` use genotypes, summary statistics and a region of interest to extract putative causal variants under a Bayesian framework with FINEMAP [21], CaviarBF [35] and PAINTOR or using stepwise model selection (cojo-slct). Output includes results of all steps and plots of regions of interest with *p*-value and post probabilities obtained by fine-mapping to compare results. In `finemapping/finemap_region.nf`, if no genotype is given by users, data are downloaded from the KGP and LD is computed (`build_example_data/main.nf`). `finemapping/cond-assoc.nf` test Independence between lead SNP and list of SNPs using GEMMA software.

#### Meta-analysis and multi-trait genome-wide association

Script `mtag-assoc` performs a multi-trait genome-wide association using the `mtag` software [23] for joint analysis of summary statistics from GWASs of different traits. The `meta-assoc.nf` workflow performs metaanalysis with different software and statistical approaches to account for variability between data sets, genomic inflation or overlap between samples with METAL [22] or GWAMA [21] and Metasoft [20,36]. Summary statistics, results of meta-analysis, and a report are produced as output.

### Format conversion

Many GWAS tools use different formats and being able to convert easily between them is useful. We provide various scripts to support this conversion. For instance the `formatdata/plk_in_vcf_imp.nf` script prepares data for imputation. There are scripts that transform VCF data imputed in various formats to PLINK, bimbam, BGEN or impute2 format. `formatdata/convert_posversiongenome.nf` converts genomic coordinates between different assemblies, for example between GRCh38 and hg19, using CrossMap [30].

### Example data set

There is a sample data set, built using KGP and GWAS catalog [37] data, at https://github.com/h3abionet/h3agwas-examples. This includes summary statistics, PLINK data, dosage, and phenotype data. For each individual in the KGP, we extracted genotype data at each position in the H3Africa Custom Array chipinfo.h3abionet.org. Data was imputed using the Sanger imputation server (https://imputation.sanger.ac.uk/). After formatting, we extracted 500 individuals and 50,000 positions.

### Installation and support

The H3AGWAS workflow requires Java 8 or later and Nextflow, and can either be cloned from GitHub explicitly or run directly using Nextflow.

In addition, the workflow relies on a number of state-of-the art bioinformatics tools (Table 1-2). We recommend that users install either Singularity or Docker and then run H3AGWAS workflow workflow using the appropriate profile – we provide containers with all tools bundled. These containers will automatically be installed on the first execution of the workflow. However, for those users who are not able to use Singularity or Docker or who would like control over which versions of the tools are used, the Docker files can be used to guide someone with basic system administration skills to install the necessary dependencies.

Manuals and examples can be found at https://github.com/h3abionet/h3agwas and https://github.com/h3abionet/h3agwas-examples. Common problems faced by users or help with the workflow itself is provided through GitHub issues. The H3ABioNet supports general queries from African researchers about the use of the workflow or GWAS in general through its help desk [38] (https://helpdesk.h3abionet.org).

### FAIR

The workflow was developed to be “Findable, Accessible, Interoperable and Reusable” according to guidelines on the FAIR https://fair-software.eu/website. The H3AGWAS workflow has been registered in bio.tools (https://bio.tools/h3agwas), uses an MIT Licence, contain citation metadata files, and uses a software quality checklist via a Core Infrastructure Initiative (CII) Best Practices badge (https://bestpractices.coreinfrastructure.org/en).

## Results and discussion

Each workflow was tested on the Wits University Core Research Cluster (CentOS 7, SLURM) and Singularity images [39], on Amazon AWS and Microsoft Azure. It has also been used in production on other environments. Since it uses Nextflow and containers, it can run on any environment that Nextflow supports such as PBS/Torque.

We illustrate the use of the workflow with a real data set from the H3Africa AWI-Gen Collaborative Centre [40]. The data comes from a cross-sectional study that investigated populations from six sub-Saharan African sites – ≈12,000 black African men and women from two urban settings (Nairobi and Soweto) and four rural settings (Agincourt, Dikgale, Nanoro and Navrongo), aged 40 to 80 years. DNA from these individuals was genotyped on the H3Africa Custom Array (https://chipinfo.h3abionet.org), designed as an African common variant enriched GWAS array with ≈2.3 million SNPs. QC was run on the array data set resulting in ≈ 10, 600 individuals and ≈1.733m SNPs. Imputation was performed on the cleaned data set using the Sanger Imputation Server and the African Genome Resources as a reference panel. We selected EAGLE2 [41] for pre-phasing and the default PBWT algorithm was used for imputation. The resulting data was used for the following phases.

### Testing of different sub-workflows

- QC : Quality control of genotype data was tested using AWI-Gen data set with 12,000 individuals before imputation.
- Association testing: For association testing, we used four residuals of lipid phenotype: LDL, cholesterol, HDL and triglycerides normalised using sex and age followed by an inverse normal transformation previously described [42]. We simultaneously ran linear associations with PLINK [18], GEMMA using the Univariate Linear Mixed Model [12], BOLT-LMM using mixed model analysis [13], fastGWA from GCTA [14, 15] using mixed linear model, SAIGE [16] and regenie [17] with genotype and dosage using BGEN format as input.
- Meta-analysis : The meta-analysis workflow was tested using GEMMA summary statistics of cholesterol from each region of AWI-Gen data set: South Africa, east Africa and west Africa.
- Other scripts : Testing of other scripts is summarized in Table 3. The `finemapping` script was tested using cholesterol result of GEMMA. Conversion of PLINK to VCF was tested using genotypes processed by the QC workflow. Conversion of VCF to PLINK, bimbam, impute2 was tested using data after imputation.

**Table 3:** List of evaluation of additional workflow implemented in H3AGWAS workflow. using AWI-Gen data set or 1000 genome project

| Script | Test descriptive |
| --- | --- |
| `qc/main.nf` | QC of genotype 12,000 individuals from AWI-Gen project |
| `finemapping/main.nf` | Extraction of lead SNPs of cholesterol result from GEMMA |
| `heritabilities/main.nf` | Estimation of heritabilities of 4 phenotypes lipid using genotype and phenotype and/or summary statistics |
| `formatdata/plk_in_vcf_imp` | Transformation of PLINK after qc in vcf to prepared data for imputation |
| `formatdata/vcf_in_plink.nf` | Transformation of data imputed in PLINK format |
| `meta/mtag-assoc.nf` | mtag performed using 4 lipid phenotypes |
| `utils/build_example_data/main.nf` | Build an example data set using diabetes phenotype from lead snps of GWAS catalog and 1000 genomes project genotype [34] |

### Association testing

The association workflow was tested using 10,700 individuals, four phenotypes and 14 million imputed positions using genotype in PLINK format and/or dosage with BGEN format [43] with PLINK, GEMMA, BOLT-LMM, fastGWA, SAIGE and regenie. We excluded Fast-LMM from testing given that it required over 100 GB of memory for a single chromosome. Using the Wits Core cluster^1^, the workflow ran with an elapsed time of 12h 36m. Among the five programs used for association, GEMMA used most computing time and jobs, followed by fastGWA, regenie, SAIGE, BOLT-LMM and PLINK. Other processes took less than 6% of CPU time Supplementary table 12. The largest maximum memory (resident set size) used by any job was 7.9 GB. Example of report of workflow can be found in supplementary text, section 3.2.

### Meta-analysis workflow

As an illustration, we performed meta-analysis (`meta/meta-assoc.nf`) using 3 summary statistics, each with 14 million SNPs. The script ran for 34 minutes in total, with METAL using the shortest processing time (1.8 minutes) and GWAMA using the longest processing time. The highest amount of memory (10 GB) was also used by GWAMA, whereas PLINK used the lowest (2 GB; Supplementary table 13).

### Others tests

Each script has been tested using the AWI-Gen data set, as summarised in Table 3. The supplementary materials provide more details, showing the costs of each step being run on a Linux cluster with SLURM and using Singularity images.

### Cloud computing

The QC and association workflows have been tested on Amazon Web Services (AWS) as well as Microsoft Azure using batch processing through Nextflow. All workflows have configuration files that include profiles for use on AWS and Azure, and instructions are provided in the README for the workflow. Using a large simulated data set with 22k individuals across 2.2m SNPs, the QC script took 8.6 hours to run on AWS and 20 hours to run on Azure, with cost between US$5-US$10 using spot pricing.

### Contribution and related work

The H3AGWAS workflow provides a comprehensive suite of portable and scalable workflows for GWAS. Few existing workflows integrate so many steps of GWAS, from QC to post-association analysis.

The closest competing workflow is BIGwas [44] which provides both QC and association testing. Kässens *et al*. compared BIGwas to an earlier version of H3AGWAS workflow. With respect to QC, they found that the two were roughly equivalent in functionality but BIGwas was much faster. However, we have been unable to replicate their findings and our experimentation shows that the QC and association testing using H3AGWAS workflow execution with default parameters is much faster (see supplementary material). However, although workflow engineering is important to performance, the computational cost primarily depends on underlying tools rather than the virtues of the workflow. With respect to association testing and preand post-analysis, they found their workflow to be superior. Whatever arguable shortcomings the H3AGWAS workflow may have had in October 2020, in March 2022 the H3AGWAS workflow has significantly more extensive set of functionalities. In addition, the H3AGWAS workflow has two significant advantages: (1) it supports cloud computing directly through the use of AWS and Azure batch; and (2) relatively lightweight Singularity/Docker containers allow deployment in HPC environments where *setuid* for Singularity is often disabled (see the supplementary for an explanation).

Other tools that are available are summarised in Table 4.

**Table 4:** Non-exhaustive list of workflows that perform QC, association testing and/or postassociation analysis of GWAS

| Software | Analysis | link |
| --- | --- | --- |
| GWASTools (Bioconductor) | QC | https://www.bioconductor.org/packages/release/bioc/html/GWASTools.html, [45] |
| BigGWAS | QC, Association testing | https://github.com/ikmb/gwas-assoc, [44] |
| plinkQC | QC | https://meyer-lab-cshl.github.io/plinkQC/articles/plinkQC.html, [46] |
| GWAS-ellingson | QC | https://github.com/sallyrose0425/GWAS, [47] |
| TASSEL | Association testing with general linear model and mixed linear model approaches, some post analysis analysis | https://avikarn.com/2019-07-22-GWAS/, [48] |
| CAUSALdb-finemapping | Fine-mapping using different software | https://github.com/mulinlab/CAUSALdb-finemapping-pip, [49, 50] |
| FINNGEN | Fine-mapping | https://github.com/FINNGEN/finemapping-pipeline |
| FUMA | post-association analysis using web interface | https://fuma.ctglab.nl/, [51, 52] |
| postgap | post-association analysis with annotation through cis-regulatory data sets using python | https://github.com/Ensembl/postgap, [53] |
| nf-gwas-pipeline | QC, Association testing integrating R packages SNPRelate/GENESIS/GMMAT and ANNOVAR using Nextflow | https://github.com/montilab/nf-gwas-pipeline [54] |
| gwasglue | post-association analysis with colocalisation, fine-mapping Mendelian randomisation using R | https://mrcieu.github.io/gwasglue |

## Conclusion

The H3AGWAS workflow provides a suite of workflows from quality control of genomic data to post-association analysis of result. Using Nextflow and containers, supports easy installation of the workflow and makes it portable and scalable – from laptop to server to cloud (AWS and Azure). The multiple workflow scripts intuitively map to individual GWAS workflow phases.

Pre-association scripts focus on quality control, with imputation performed by a separate workflow. We plan to add calling of array data to the workflow, in the future. Association studies, including G *×* E analysis, can be performed in our workflow using six different techniques provided by state-of-the-art tools. Post-analysis of GWAS supports meta analysis, heritability computation, identifying causal SNPs, co-localisation and fine-mapping.

Our workflow supports multiple tools, providing users with opportunities to compare results (e.g., different approaches for fine-mapping and association testing). Furthermore, different Nextflow scripts for each step allows the user to run analyses with different parameters and customise the analysis to their needs. Each script is associated with a Docker image to simplify installation, and returns a PDF report to the researcher to help to interpret the results.

The workflows are available on GitHub and we strive to comply with FAIR principles.

### Future development

Several additional features are under development. In pre-association, calling genotypes from raw array data is challenging, and we are currently working on a workflow to perform this step. New features to be added include supporting replication and transferability of previous result using GWAS Catalog result [37] or full summary statistics. We also plan to port the workflow to DSL2 and make it nf-core compatible.

## Availability and requirements

|  |  |
| --- | --- |
| **Project name:** | H3AGWAS workflow |
| **Project home page:** | https://github.com/h3abionet/h3agwas |
| **Example home page:** | https://github.com/h3abionet/h3agwas-example |
| **Operating system:** | Linux (or MacOS and Windows with Docker) |
| **Program language:** | Nextflow, Python, R, bash |
| **Other requirements:** | Java 8 or later, Nextflow<br>Docker/Singularity (or software dependencies listed in Dockerfile) |
| **Licence:** | MIT Licence |
| **Restrictions on non-academic use:** | None |

Docker images are available from https://quay.io/organization/h3abionet_org/ and https://github.com/h3abionet/h3agwas-docker.

Example are available from https://github.com/h3abionet/h3agwas-examples.

### Ethics

We have used the AWI-Gen data set as our main example as a real data set. The AWI-Gen study received approval from the Human Research Ethics Committee (Medical), University of the Witwatersrand, South Africa (M121029, M1706110). The AWI-Gen data set is available from the European Genome-Phenome Archive on application to the independent H3Africa Data Access Committee (EGAD00001006425 and EGAD00010001996). We hope to make our synthetic data sets publicly available.

### Funding

The work is supported by National Human Genome Research Institute/National Institutes of Health: JTB is supported by the AWI-Gen Collaborative Centre (U54HG006938) and all other authors and the Wits Core Cluster are supported by the Pan-African Bioinformatics Network for H3Africa (U24HG006941). The views expressed are solely those of the authors and not that of the NIH.

### Conflict interest and consents

Each authors contribute and corrected papers and consent for publication. There is no conflict of interest.

### Authors contribution

JTB developed the workflow and led the writing of this paper. LC contributed to worflow development. SB, CJF and SP led the H3ABioNet workflows project and provided scientific input and direction. SH led the H3AGWAS workflow project, and wrote parts of the workflow, and co-led the writing of the paper. All authors contributed to the writing of this paper.

## Acknowledgements

The authors thank all AWI-Gen collaborators for use of their data set and more particularly Michèle Ramsay Principal Investigator and all participants of AWI-Gen. Our SBIMB colleagues have made many useful and generous contributions: Carl Wenlong Chen, Palwendé R. Boua, Vivien Chebii and Shaun Aron in particular. Many people contributed to the workflow and we especially thank Lerato Magosi, Rob Clucas and Eugene de Beste whose effort at the start of the project was so important. We thank Professor Nicola Mulder from the University of Cape Town whose leadership of H3ABioNet made the work possible.

