## supplementary for "H3AGWAS : A portable workflow for Genome Wide Association Studies"

Brandenburg *et al*

August 8, 2022

**4 Detail of experimentation in the main paper 27**

### 1 Overview

This supplementary note contains two substantive sections: The first give a comparison between H3AGWAS workflow and a competing tool, especially with respect to computational performance.

The second gives example runs of a number of the different workflows that we provide.

### 2 Computational comparison between the BIGwas and H3AGWAS workflow

#### 2.1 Introduction

This section documents a computational comparison of the BIGwas and H3AGWAS workflow workflows in response to the findings of Kässens *et al.* that BIGwas is significantly faster than H3AGWAS workflow for QC for medium and larger files. For example, Kässens *et al.* found that data set with 5k individuals and 50k SNPs takes 8 minutes (BIGwas) versus 15 minutes (H3AGWAS workflow), and a data set with 20,554 individuals and 700k SNPs takes 135m (BIGwas) versus 537m (H3AGWAS workflow) and for even larger data sets that they could not reasonably complete execution of H3AGWAS workflow. (It must be emphasised that comparing workflows requires comparing multiple factors and in our view this is not the most important factor, but Kässens *et al* have made very serious negative findings, which we believe needs to be addressed. We do not agree with their findings)

#### 2.2 Data sets used

The following data sets were used:

- The example data set that comes with BIGwas – this is a set based on 1000 Genomes data with 2504 individuals and 50k SNPs.
- The AWI-Gen unqc-ed data – 11062 individuals and 2.267 million SNPs.
- A simulated data set (*sim1* 22142 individuals and 2.267 million SNPs.

One difference between the two workflows is that H3AGWAS workflow expects that all the genotype data is in one PLINK file, while BIGwas allows multiple files to be input which are then merged, which exposes further parallelism. To make the comparison fair we compare the H3AGWAS workflow times with two separate runs of the BIGwas workflow:

- The BIGwas example data set is provided as two separate PLINK data sets – one the "cases" and one the "controls", each with approximately 2500 individuals. We compare the H3AGWAS workflow with the merged data setas input with the BIGwas on both the merged data and on the original split files.
- For the AWI-Gen data, we split into roughly two halves and artificially declare the two halves as cases and controls. We compare the H3AGWAS workflow on the original AWI-Gen data with BIGwas on the original data and on the split data. For the *sim1* data set a similar comparison was made.

#### 2.3 Computational setup

We used Nextflow 21.04.1 and used Singularity with the images provided by the two tools. In both cases, both the Nextflow repos and the Singularity images were already downloaded and installed. We performed the experiments on a single machine (no scheduler) and using SLURM on a cluster.

##### 2.3.1 Single machine execution

We used a machine with a dual Xeon Silver 4214 CPUs running at 2.20GHz (24 physical cores, 48 hyper-threaded cores) and 128GB of RAM, and all data inputs and outputs were stored on a Seagate ST2000NM0008 2TB SATA Hard Drive. The machine was otherwise unloaded.

We ran both of these on the default settings – presumably both developers chose appropriate default settings so that is fair enough. However, H3AGWAS workflow has performance parameters that the user can set if they have more powerful computers. The `max_plink_cores` parameter is

| Data set | H3AGWAS workflow | | BIGwas | | | |
|---|---|---|---|---|---|---|
| | | | Split | | Merged | |
| | Elapsed (s) | CPU h | Elapsed (s) | CPU h | Elapsed (s) | CPU h |
| Example | 34 | 0.05 | 480 | 0.1 | 496 | 0.2 |
| AWI-Gen | 1383 | 2.1 | 2576 | 0.8 | 3700 | 1.2 |
| sim1 | 7278 | 12.3 | 29938 | 8.3 | 30240 | 8.4 |

Table 1: *Comparison between H3A*GWAS *workflow and BIGwas workflow using QC script and 3 different data sets using single machine execution*

by default set to 4 – this limits any PLINK process to use 4 cores (and makes 4 cores available and so hence counts to CPU hours whether used or not). If we run the Nextflow with the `--max_plink_cores=12`, the elapsed time for the AWI-Gen data set drops from 1383s to 750s at the cost of accounted CPU hours going to 4.2 CPU hours. This is a trade-off the user must consider. No doubt there are similar changes that could be made to the BIGwas workflow.

Note the difference in the AWI-Gen and *sim1* data set for the H3AGWAS workflow is a factor of 5.2. In principle we would expect the overall computational cost to scale quadratically with number of SNPs as the single biggest computational costs are steps which are quadratic. However as these components take longer there is less task parallelism as a proportion of the overall cost. The BIGwas workflow also scales super-quadratically but by a greater factor (our superficial observation is that the bulk of this extra cost is as at similar points in the computation).

##### 2.3.2 Cluster execution

We tested on our production University Research Cluster: SLURM 20.11.8, CentOS7.9, Singularity 3.6.3 (default Singularity OSG release). The cluster is heterogeneous so we set Nextflow *clusterOptions* to execute only on 20 nodes with dual core Intel Xeon Silver 4114 CPUs running at 2.2GHz (20 physical, 40 hyper-threaded cores per node – note that these machines are slower than the one we tested above). Since this is a production cluster we were unable to test while the cluster otherwise completely idle but we could test late on weekend with only a few other jobs running so we do not think that this affected the results (we manually inspected that the jobs ran on machines not being used by other jobs).

**Singularity issues:** We were unable to run BIGwas on the cluster in its default Singularity setting. Like many production HPC systems, the cluster follows recommendations to not allow Singularity with *setuid* enabled (`https://sylabs.io/guides/latest/user-guide/security.html`). The disadvantage of this is that standard Singularity SIF images cannot be directly executed but must be copied (unsquashed) to a temporary disk for each separate Nextflow process[1]. The BIGwas image is 11GB in size in SIF (compressed) format and the computational cost of unsquashing, especially when multiple processes are doing this in parallel made running the workflow impractical. The same Singularity issue applies to H3AGWAS workflow, of course. However, the H3AGWAS workflow container design approach is to have several, specialised containers rather than one monolithic container. For example, the workhorse `py3plink` container is ≈450MB in size – so although there is a significant penalty for running H3AGWAS workflow on the cluster using Singularity compared to running the natively installed software, it is not an outrageous penalty. In order to perform the testing below, we were able to enable setuid for (this requires root privileges), but this is not something we are allowed to leave for extended periods. In many environments setuid is enabled in Singularity installations (as it was in our testing on the single machine), so many users will not face this problem. But we suspect that other Singularity users in production HPC environment will run into the same problem as us. (We also tested Singularity 3.8 and 3.9 and it had the same problem).

[1]And to be clear if a Nextflow process is executed 10 times in parallel then each instantiation of the process requires this.

| Data set | H3AGWAS workflow | | BIGwas | | | |
|---|---|---|---|---|---|---|
| | | | Split | | Merged | |
| | Elapsed (s) | CPU h | Elapsed (s) | CPU h | Elapsed (s) | CPU h |
| Example | 189 | $< 0.1$ | 967 | 0.4 | 899 | 0.2 |
| AWI-Gen | 2025 | 3.8 | 3712 | 1.2 | 3647 | 1.1 |
| sim1 | 8346 | 16.2 | 33728 | 8.4 | 31028 | 8.6 |

Table 2: *Comparison between H3A*GWAS *workflow and BIGwas workflow using the QC script and three different data sets using the Cluster and SLURM*

#### 2.4 Comparison between association workflow of BIGwas and H3AGWAS workflow

##### 2.4.1 Data sets used and methodology

We used output of QC produced by the BIGwas workflow to run association of H3AGWAS workflow and BIGwas. For both workflow we ran PLINK as the underlying association testing tool defaults of workflows where used. We ran each test using an Intel Xeon Silver 4114 dual core processor (40 hyper-threaded cores on 20 physical cores) with 128GB of RAM

#### 2.5 Run, troubleshooting and duration

Table 3 shows the comparison. For the small test Data set, duration is lower for BIGwas than H3AGWAS workflow, but for bigger sample size and SNP number, H3AGWAS workflow performed better.

In using BIGwas, we observed missing data causes error on workflow when used with binary phenotype.

| Data set | SNPs number | Sample Size | BIGwas | H3AGWAS workflow |
|---|---|---|---|---|
| Example | 35317 | 2464 | 35 | 70 |
| AWI-GEN | 2120006 | 8487 | 2383 | 1637 |
| sim1 | 2091083 | 18322 | 5607 | 2327 |

Table 3: Comparison between H3AGWAS workflow and BIGwas workflow using Association script and 3 different data sets

#### 2.6 Conclusion

As we have indicated, performance is not the primary measure of a workflow because ultimately the costs depend on the underlying software used and the workflow designer can neither take too much credit nor blame for this. However, it is important to demonstrate the workflow can expose appropriate parallelism, which we believe has been demonstrated. Certainly we do not believe that the experimental evidence supports the claim that the BIGwas workflow is faster than H3AGWAS workflow.

### 3 Description and test of different scripts of H3AGWAS workflow : CPUs, Time

This section shows additional testing of H3AGWAS workflow on our SLURM cluster as shown. Again since the cluster is a production cluster we were only able to run it on a lightly loaded cluster not on a cluster that was idle. The purpose of this testing is to give indicative real-world costs. For the tests done, the workflow execution is shown graphically as a directed acyclic graph and the computational cost of the individual components is shown (of course, many of the individual components can be done in parallel).

#### 3.1 Quality Control of genetics data

- Objectives : apply a quality control on genetics data.
- Input : workflow take as input PLINK file from genomics data, phenotype, sex phenotype.
- Individual filter :
  - Apply sex control with X chromosome and sex phenotype.
  - heterozygosity control using Hardy–Weinberg equilibrium.
  - missingness
  - relatedness
- SNPs filter :
  - minor allele frequency
  - heterozygosity
  - duplicated markers
  - missingness
- Output :
  - report in pdf is produce in PDF describing each steps with different filters
  - PLINK file after quality control with frequencies distribution, hardy Weinberg equilibrium... see example 1
  - intermediate files produce by workflow.
- Test: QC workflow has been apply for 12,000 individuals with genotype using h3array positions (2.4 millions positions) using the cluster and SLURM – see the statistics in table 4.

An overview of the execution can be found in Figure 2 and the detailed computational results in Table 4.

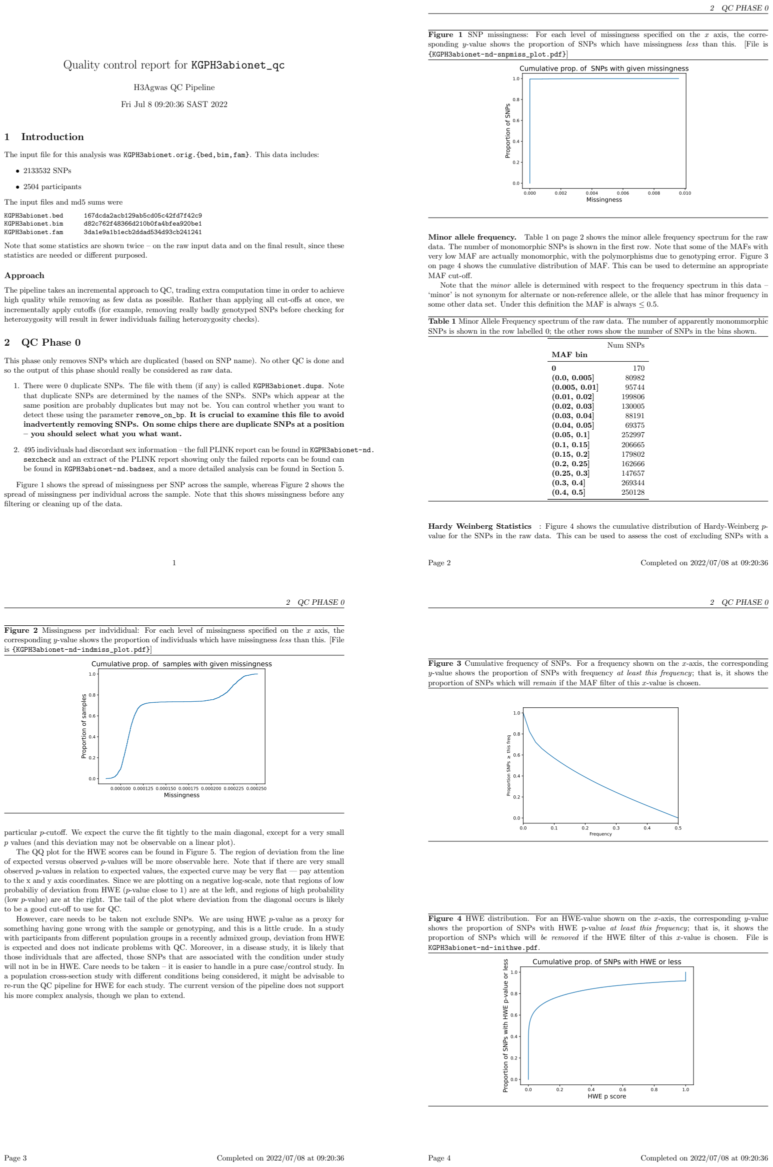

Quality control report for `KGPH3abionet_qc`

H3Agwas QC Pipeline

Fri Jul 8 09:20:36 SAST 2022

#### 1 Introduction

The input file for this analysis was `KGPH3abionet.orig.{bed,bim,fam}`. This data includes:

- 2133532 SNPs
- 2504 participants

The input files and md5 sums were

```
KGPH3abionet.bed     167dcda2acb129ab5cd05c42fd7f42c9
KGPH3abionet.bim     d82c762f48366d210b0fa4bfea920be1
KGPH3abionet.fam     3da1e9a1b1ecb2ddad534d93cb241241
```

Note that some statistics are shown twice – on the raw input data and on the final result, since these statistics are needed or different purposed.

##### Approach

The pipeline takes an incremental approach to QC, trading extra computation time in order to achieve high quality while removing as few data as possible. Rather than applying all cut-offs at once, we incrementally apply cutoffs (for example, removing really badly genotyped SNPs before checking for heterozygosity will result in fewer individuals failing heterozygosity checks).

#### 2 QC Phase 0

This phase only removes SNPs which are duplicated (based on SNP name). No other QC is done and so the output of this phase should really be considered as raw data.

1. There were 0 duplicate SNPs. The file with them (if any) is called `KGPH3abionet.dups`. Note that duplicate SNPs are determined by the names of the SNPs. SNPs which appear at the same position are probably duplicates but may not be. You can control whether you want to detect these using the parameter `remove_on_bp`. **It is crucial to examine this file to avoid inadvertently removing SNPs. On some chips there are duplicate SNPs at a position – you should select what you what want.**
2. 495 individuals had discordant sex information – the full PLINK report can be found in `KGPH3abionet-nd.sexcheck` and an extract of the PLINK report showing only the failed reports can be found can be found in `KGPH3abionet-nd.badsex`, and a more detailed analysis can be found in Section 5.

Figure 1 shows the spread of missingness per SNP across the sample, whereas Figure 2 shows the spread of missingness per individual across the sample. Note that this shows missingness before any filtering or cleaning up of the data.

1

*2 QC PHASE 0*

**Figure 1** SNP missingness: For each level of missingness specified on the $x$ axis, the corresponding $y$-value shows the proportion of SNPs which have missingness *less* than this. [File is `{KGPH3abionet-nd-snpmiss_plot.pdf}`]

**Minor allele frequency.** Table 1 on page 2 shows the minor allele frequency spectrum for the raw data. The number of monomorphic SNPs is shown in the first row. Note that some of the MAFs with very low MAF are actually monomorphic, with the polymorphisms due to genotyping error. Figure 3 on page 4 shows the cumulative distribution of MAF. This can be used to determine an appropriate MAF cut-off.

Note that the *minor* allele is determined with respect to the frequency spectrum in this data – 'minor' is not synonym for alternate or non-reference allele, or the allele that has minor frequency in some other data set. Under this definition the MAF is always $\leq 0.5$.

**Table 1** Minor Allele Frequency spectrum of the raw data. The number of apparently monommorphic SNPs is shown in the row labelled 0; the other rows show the number of SNPs in the bins shown.

| MAF bin | Num SNPs |
|---|---|
| **0** | 170 |
| **(0.0, 0.005]** | 80982 |
| **(0.005, 0.01]** | 95744 |
| **(0.01, 0.02]** | 199806 |
| **(0.02, 0.03]** | 130005 |
| **(0.03, 0.04]** | 88191 |
| **(0.04, 0.05]** | 69375 |
| **(0.05, 0.1]** | 252997 |
| **(0.1, 0.15]** | 206665 |
| **(0.15, 0.2]** | 179802 |
| **(0.2, 0.25]** | 162666 |
| **(0.25, 0.3]** | 147657 |
| **(0.3, 0.4]** | 269344 |
| **(0.4, 0.5]** | 250128 |

**Hardy Weinberg Statistics** : Figure 4 shows the cumulative distribution of Hardy-Weinberg $p$-value for the SNPs in the raw data. This can be used to assess the cost of excluding SNPs with a

Page 2 Completed on 2022/07/08 at 09:20:36

*2 QC PHASE 0*

**Figure 2** Missingness per indvididual: For each level of missingness specified on the $x$ axis, the corresponding $y$-value shows the proportion of individuals which have missingness *less* than this. [File is `{KGPH3abionet-nd-indmiss_plot.pdf}`]

particular $p$-cutoff. We expect the curve the fit tightly to the main diagonal, except for a very small $p$ values (and this deviation may not be observable on a linear plot).

The QQ plot for the HWE scores can be found in Figure 5. The region of deviation from the line of expected versus observed $p$-values will be more observable here. Note that if there are very small observed $p$-values in relation to expected values, the expected curve may be very flat — pay attention to the x and y axis coordinates. Since we are plotting on a negative log-scale, note that regions of low probabiliy of deviation from HWE ($p$-value close to 1) are at the left, and regions of high probability (low $p$-value) are at the right. The tail of the plot where deviation from the diagonal occurs is likely to be a good cut-off to use for QC.

However, care needs to be taken not exclude SNPs. We are using HWE $p$-value as a proxy for something having gone wrong with the sample or genotyping, and this is a little crude. In a study with participants from different population groups in a recently admixed group, deviation from HWE is expected and does not indicate problems with QC. Moreover, in a disease study, it is likely that those individuals that are affected, those SNPs that are associated with the condition under study will not in be in HWE. Care needs to be taken – it is easier to handle in a pure case/control study. In a population cross-section study with different conditions being considered, it might be advisable to re-run the QC pipeline for HWE for each study. The current version of the pipeline does not support his more complex analysis, though we plan to extend.

Page 3 Completed on 2022/07/08 at 09:20:36

*2 QC PHASE 0*

**Figure 3** Cumulative frequency of SNPs. For a frequency shown on the $x$-axis, the corresponding $y$-value shows the proportion of SNPs with frequency *at least this frequency*; that is, it shows the proportion of SNPs which will *remain* if the MAF filter of this $x$-value is chosen.

**Figure 4** HWE distribution. For an HWE-value shown on the $x$-axis, the corresponding $y$-value shows the proportion of SNPs with HWE p-value *at least this frequency*; that is, it shows the proportion of SNPs which will *be removed* if the HWE filter of this $x$-value is chosen. File is `KGPH3abionet-nd-inithwe.pdf`.

Page 4 Completed on 2022/07/08 at 09:20:36

Figure 1: Example of four first pages of quality control report generated by pipeline

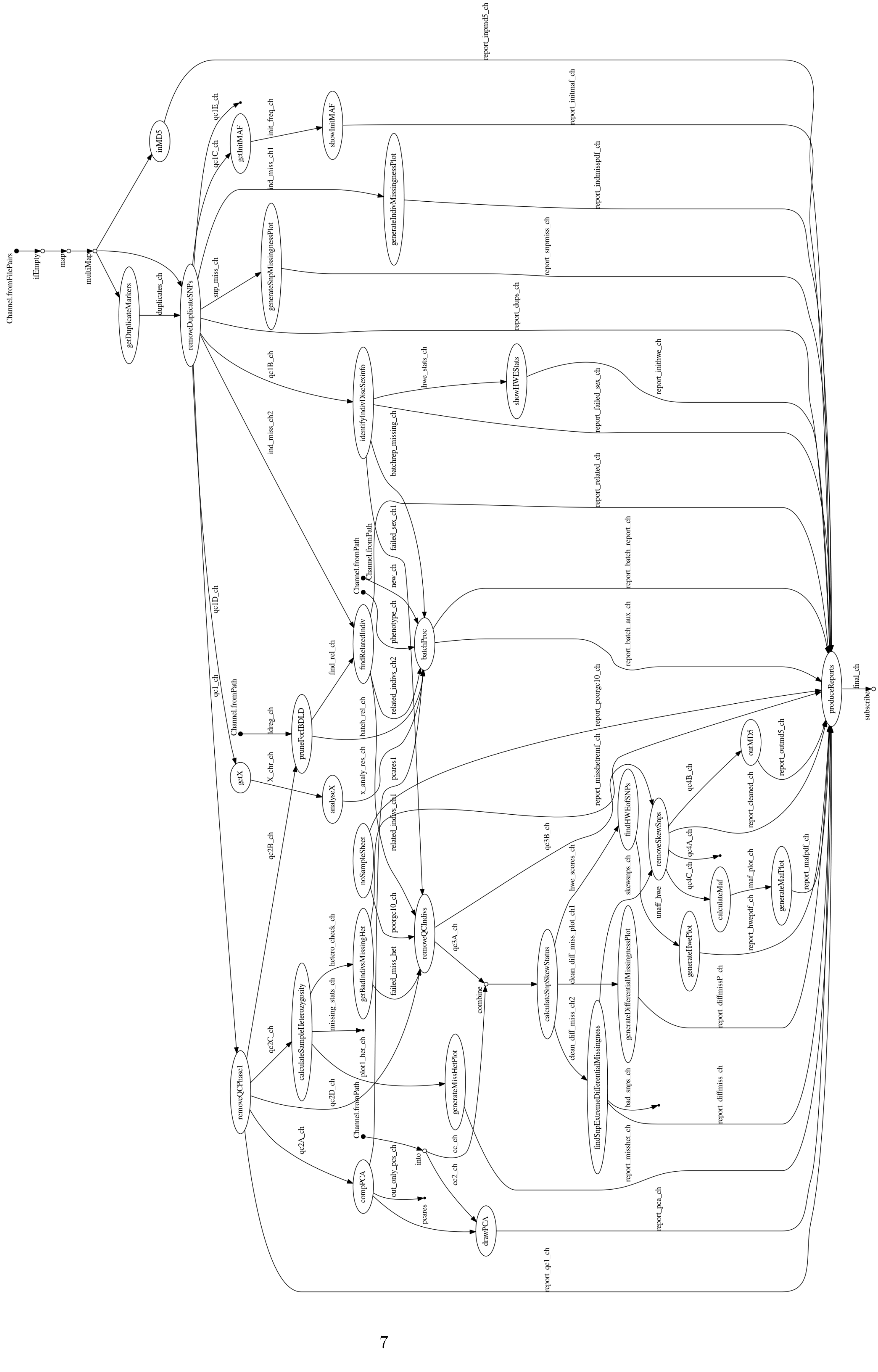
Channel.fromFilePairs
ifEmpty
map
multiMap
inMD5
report_inpmd5_ch
getDuplicateMarkers
duplicates_ch
removeDuplicateSNPs
qc1E_ch
qc1C_ch
getInitMAF
init_freq_ch
showInitMAF
report_initmaf_ch
ind_miss_ch1
generateIndivMissingnessPlot
report_indmisspdf_ch
snp_miss_ch
generateSnpMissingnessPlot
report_snpmiss_ch
report_dups_ch
qc1B_ch
identifyIndivDiscSexinfo
hwe_stats_ch
showHWEStats
report_inithwe_ch
report_failed_sex_ch
batchrep_missing_ch
ind_miss_ch2
report_related_ch
failed_sex_ch1
Channel.fromPath
Channel.fromPath
new_ch
phenotype_ch
report_batch_report_ch
batchProc
report_batch_aux_ch
findRelatedIndiv
find_rel_ch
related_indivs_ch2
qc1D_ch
qc1_ch
Channel.fromPath
idreg_ch
pruneForIBDLD
batch_rel_ch
report_poorgc10_ch
getX
X_chr_ch
analyseX
x_analy_res_ch
pcares1
report_misshetremf_ch
related_indivs_ch1
noSampleSheet
qc3B_ch
findHWEofSNPs
removeSkewSnps
qc4B_ch
outMD5
report_outmd5_ch
report_cleaned_ch
qc2B_ch
hetero_check_ch
poorgc10_ch
removeQCIndivs
hwe_scores_ch
skewsnps_ch
qc4A_ch
qc4C_ch
calculateMaf
maf_plot_ch
generateMafPlot
report_mafpdf_ch
unaff_hwe
generateHwePlot
report_hwepdf_ch
getBadIndivsMissingHet
failed_miss_het
qc3A_ch
clean_diff_miss_plot_ch1
calculateSnpSkewStatus
generateDifferentialMissingnessPlot
report_diffmissP_ch
calculateSampleHeterozygosity
missing_stats_ch
combine
clean_diff_miss_ch2
findSnpExtremeDifferentialMissingness
plot1_het_ch
qc2C_ch
generateMissHetPlot
qc2D_ch
bad_snps_ch
report_diffmiss_ch
removeQCPhase1
qc2A_ch
Channel.fromPath
report_misshet_ch
into
cc_ch
out_only_pcs_ch
compPCA
cc2_ch
pcares
drawPCA
report_pca_ch
report_qc1_ch
produceReports
final_ch
subscribe

| Process | Tot hours | % times | % cpu number used (Mean) | Max mem (MB) | NF processes |
|---|---|---|---|---|---|
| analyseX | 0.00 | 1.13 | 28.40 | 67.10 | 1 |
| batchProc | 0.00 | 0.31 | 106.50 | 100.70 | 1 |
| calculateMaf | 0.00 | 0.23 | 55.10 | 138.20 | 1 |
| calculateSampleHeterozygosity | 0.10 | 2.40 | 25.70 | 105.90 | 1 |
| calculateSnpSkewStatus | 0.00 | 1.03 | 295.90 | 212.60 | 1 |
| compPCA | 0.60 | 19.73 | 365.40 | 1200.00 | 1 |
| drawPCA | 0.00 | 0.54 | 85.70 | 83.60 | 1 |
| findHWEofSNPs | 0.00 | 0.98 | 5.30 | 2.40 | 1 |
| findRelatedIndiv | 0.00 | 0.40 | 32.80 | 49.70 | 1 |
| findSnpExtremeDifferentialMissingness | 0.00 | 0.33 | 101.30 | 125.50 | 1 |
| generateDifferentialMissingnessPlot | 0.00 | 0.33 | 107.60 | 327.70 | 1 |
| generateHwePlot | 0.00 | 0.91 | 54.30 | 344.20 | 1 |
| generateIndivMissingnessPlot | 0.00 | 0.33 | 136.20 | 11.40 | 1 |
| generateMafPlot | 0.00 | 0.42 | 76.20 | 512.40 | 1 |
| generateMissHetPlot | 0.00 | 1.07 | 90.50 | 101.90 | 1 |
| generateSnpMissingnessPlot | 0.10 | 2.54 | 57.10 | 299.00 | 1 |
| getBadIndivsMissingHet | 0.00 | 1.41 | 61.70 | 49.70 | 1 |
| getDuplicateMarkers | 0.00 | 1.64 | 82.40 | 221.60 | 1 |
| getInitMAF | 0.10 | 2.55 | 10.80 | 171.80 | 1 |
| getX | 0.10 | 2.40 | 17.30 | 113.90 | 1 |
| identifyIndivDiscSexinfo | 0.10 | 3.33 | 25.70 | 214.90 | 1 |
| inMD5 | 0.00 | 1.64 | 23.10 | 8.10 | 1 |
| noSampleSheet | 0.00 | 1.30 | 34.10 | 61.80 | 1 |
| outMD5 | 0.00 | 0.95 | 35.30 | 8.10 | 1 |
| produceReports | 0.00 | 1.37 | 4.60 | 25.10 | 1 |
| pruneForIBDLD | 0.90 | 31.43 | 384.50 | 1500.00 | 1 |
| removeDuplicateSNPs | 0.10 | 4.17 | 15.30 | 199.50 | 1 |
| removeQCIndivs | 0.00 | 1.60 | 28.20 | 152.20 | 1 |
| removeQCPhase1 | 0.30 | 10.20 | 28.00 | 196.30 | 1 |
| removeSkewSnps | 0.00 | 0.69 | 47.80 | 152.40 | 1 |
| showHWEStats | 0.00 | 1.01 | 76.70 | 926.10 | 1 |
| showInitMAF | 0.00 | 1.62 | 51.40 | 484.80 | 1 |

Table 4: Statistics resumé of the QC workflow: process – Nextflow process name; Tot. hours – total hours used by NF process; % times – percentage of times used by process compared to other process ; % cpu number used (Mean) – mean % cpu number used by the process; Max mem (MB) is maximum of memory (resident set size) used by one process; NF processes – number of Nextflow process used for the steps

##### 3.2 Association

- input :
  - phenotype file and one or more phenotype, covariates
  - genetics data in plink file and in option dosage : bgen (regenie, SAIGE, fastGWA and BOLT-LMM), VCF (SAIGE) and impute2 (BOLT-LMM)
- output :
  - each summary statistics of each software and phenotype used and pdf report contains for each combination 3
  - report with Manhattan, qq plot and best result
  - relatedness comptued for each software

##### 3.3 Fine-mapping

- Objective: apply a fine-mapping on significant regions of summary statistics result
- Input data : summary statistics, causal variant number and genetics data in PLINK format
- identify region to apply fine-mapping on full summary statistics using PLINK clump to identify lead SNPs.
- Steps
  - for each region apply different algorithms to find the number of independent SNPs, putative causal variant and credible interval with different software for fine-mapping : COJO (step-wise model selection procedure to select independently associated), FINEMAP (stochastic and conditional algorithm), Caviarbf and PAINTOR software with possibility to use eQTL information.
- Output 4:
  - intermediate file produce in workflow and by each software
  - file contains all result merge
  - figures plot as locus zoom with has been had probability of each software of fine-mapping.
- Test : Fine-mapping has been done on summary statistics obtained with cholesterol output of association testing done with GEMMA with AWI-Gen data set. We obtained 40 windows with $p < 5 \times 10^{-8}$: for each region different software was used fine-mapping

Figure 5 gives an overview of the execution and Table 5 shows the detailed computational costs.

| Process | Tot hours | % times | % cpu number used (Mean) | Max mem (MB) | NF processes |
|---|---|---|---|---|---|
| clump_data | 0.20 | 2.19 | 61.60 | 3900.00 | 1 |
| ComputedCaviarBF | 0.50 | 5.79 | 86.80 | 16.50 | 40 |
| ComputedCojo | 0.50 | 5.54 | 76.41 | 11.10 | 40 |
| ComputedFineMapCond | 0.50 | 6.72 | 81.09 | 11.20 | 40 |
| ComputedFineMapSSS | 0.40 | 5.53 | 164.14 | 8.20 | 40 |
| ComputedLd | 0.90 | 11.41 | 113.56 | 38.40 | 40 |
| ComputedPaintor | 0.40 | 5.29 | 85.59 | 76.40 | 40 |
| extract_sigpos | 0.00 | 0.12 | 131.90 | | 1 |
| ExtractPositionGWAS | 1.90 | 22.84 | 95.63 | 3800.00 | 40 |
| GetGenesInfo | 0.10 | 1.23 | 42.60 | 2000.00 | 1 |
| GWASCatDl | 0.00 | 0.25 | 77.30 | 445.20 | 1 |
| MergeResult | 0.70 | 8.12 | 93.01 | 439.10 | 40 |
| SubPLINK | 2.00 | 25.00 | 78.50 | 826.50 | 40 |

Table 5: Statistics resumé of fine-mapping workflow running on cluster, using cholesterol phenotype. Process - it is Nextflow process name; Tot. hours – total hours used by NF process; % times – percentage of times used by process compared to other process ; % cpu number used (Mean) - mean % cpu number used by the process; Max mem (MB) is maximum of memory used by one process; NF processes - number of Nextflow process used for the steps

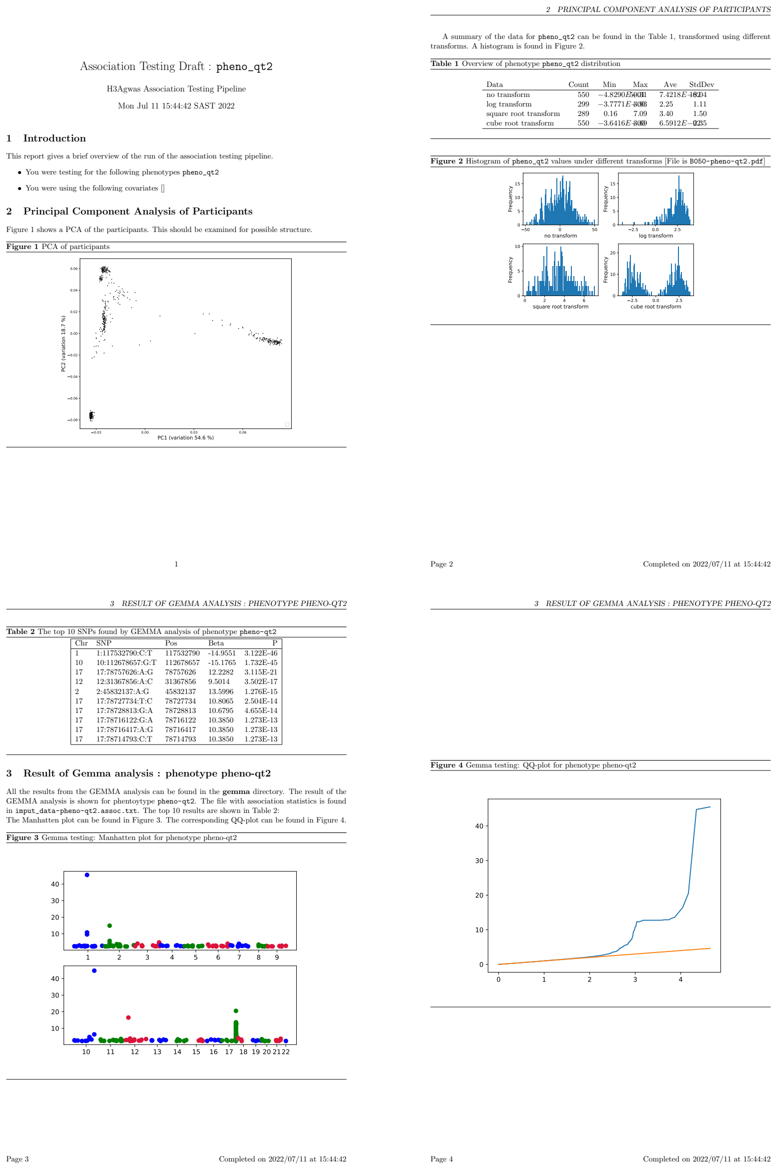

Association Testing Draft : pheno_qt2

H3Agwas Association Testing Pipeline

Mon Jul 11 15:44:42 SAST 2022

1 Introduction

This report gives a brief overview of the run of the association testing pipeline.

- You were testing for the following phenotypes pheno_qt2
- You were using the following covariates []

2 Principal Component Analysis of Participants

Figure 1 shows a PCA of the participants. This should be examined for possible structure.

**Figure 1** PCA of participants

1

2 PRINCIPAL COMPONENT ANALYSIS OF PARTICIPANTS

A summary of the data for pheno_qt2 can be found in the Table 1, transformed using different transforms. A histogram is found in Figure 2.

**Table 1** Overview of phenotype pheno_qt2 distribution

| Data | Count | Min | Max | Ave | StdDev |
|---|---|---|---|---|---|
| no transform | 550 | −4.8290E5001 (overprinted) | | 7.4218E−1804 (overprinted) | |
| log transform | 299 | −3.7771E-3.03 (overprinted) | | 2.25 | 1.11 |
| square root transform | 289 | 0.16 | 7.09 | 3.40 | 1.50 |
| cube root transform | 550 | −3.6416E-3.69 (overprinted) | | 6.5912E−0235 (overprinted) | |

**Figure 2** Histogram of pheno_qt2 values under different transforms [File is B050-pheno-qt2.pdf]

Page 2 Completed on 2022/07/11 at 15:44:42

3 RESULT OF GEMMA ANALYSIS : PHENOTYPE PHENO-QT2

**Table 2** The top 10 SNPs found by GEMMA analysis of phenotype pheno-qt2

| Chr | SNP | Pos | Beta | P |
|---|---|---|---|---|
| 1 | 1:117532790:C:T | 117532790 | -14.9551 | 3.122E-46 |
| 10 | 10:112678657:G:T | 112678657 | -15.1765 | 1.732E-45 |
| 17 | 17:78757626:A:G | 78757626 | 12.2282 | 3.115E-21 |
| 12 | 12:31367856:A:C | 31367856 | 9.5014 | 3.502E-17 |
| 2 | 2:45832137:A:G | 45832137 | 13.5996 | 1.276E-15 |
| 17 | 17:78727734:T:C | 78727734 | 10.8065 | 2.504E-14 |
| 17 | 17:78728813:G:A | 78728813 | 10.6795 | 4.655E-14 |
| 17 | 17:78716122:G:A | 78716122 | 10.3850 | 1.273E-13 |
| 17 | 17:78716417:A:G | 78716417 | 10.3850 | 1.273E-13 |
| 17 | 17:78714793:C:T | 78714793 | 10.3850 | 1.273E-13 |

3 Result of Gemma analysis : phenotype pheno-qt2

All the results from the GEMMA analysis can be found in the **gemma** directory. The result of the GEMMA analysis is shown for phentoytype pheno-qt2. The file with association statistics is found in imput_data-pheno-qt2.assoc.txt. The top 10 results are shown in Table 2:
The Manhatten plot can be found in Figure 3. The corresponding QQ-plot can be found in Figure 4.

**Figure 3** Gemma testing: Manhatten plot for phenotype pheno-qt2

Page 3 Completed on 2022/07/11 at 15:44:42

3 RESULT OF GEMMA ANALYSIS : PHENOTYPE PHENO-QT2

**Figure 4** Gemma testing: QQ-plot for phenotype pheno-qt2

Page 4 Completed on 2022/07/11 at 15:44:42

Figure 3: Example of four first pages of association report generated by pipeline with 10 best solutions, qq plot and manhantan plot

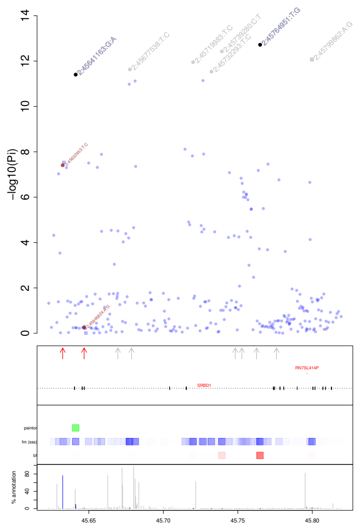

Legends (from top to bottow):

– top box:

– gwas catalog, are square point and rsid red

– lead snps defined by gcta are points and rsid in black

– credible set defined by cond / sss finemapping are in grey

– genes / gwas catalog box :

– arrow : red arrow : present in dataset / grey arrow absent in dataset

– genes represented by box (exon) and lines (intron)

– distribution of post probability :

– dstribution of post probability for finemapinf caviar, paintor and finemap, color variabtio

– distribution of annotation : in % for each positions, red correspond to lead position, or c

Gwas catalog :

45775995 Haemorrhoidal disease /PubmedId: 33888516 –

45762607 Osteoarthritis (self−reported) /PubmedId: 29559693 –

45752709 Metabolite levels /PubmedId: 23823483 –

45748277 Type 2 diabetes /PubmedId: 25760438 –

45678606 Carotid Intima−media thickness (mean of the maximum cIMT) /Pubm

45669485 Height /PubmedId: 30595370 –

45646824 Glaucoma /PubmedId: 20363506 –

45632453 Metabolite levels /PubmedId: 23823483 –

Figure 4: Example of report generated by fine-mapping pipeline contained page 1)locus-zoom with lead snps defined using stepwise model selection procedure (gcta), credible position from fine-mapping softwares, post-probability of fine-mapping software, information relative to GWAS catalog, genes. Pages 2 : legends. Pages 3 : GWAS catalog information's.

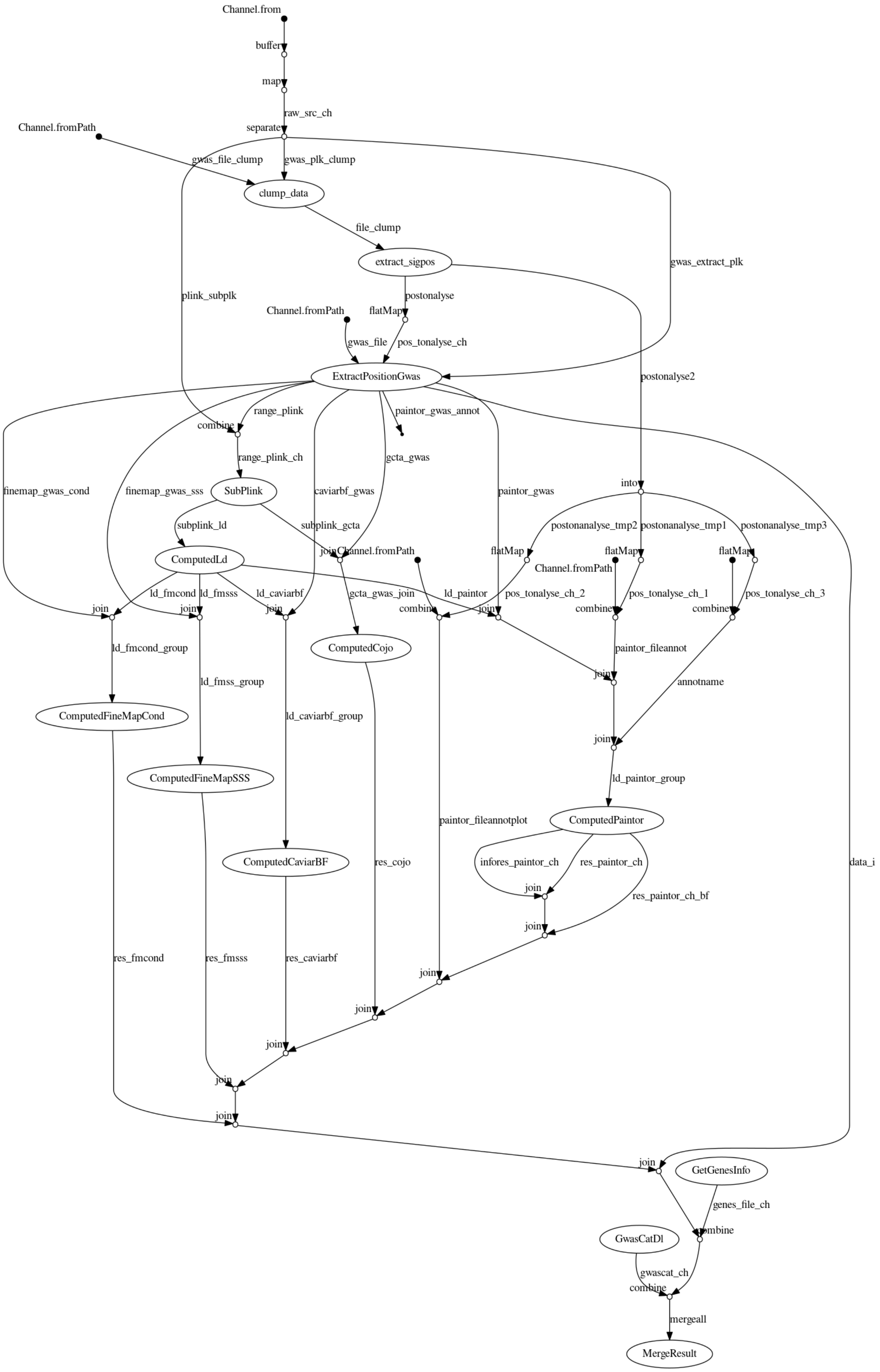

Figure 5: flowchart of Fine-mapping workflow

#### 3.4 Heritability

- Objectives : compute heritability and/or co-heritability using genetics diversity and phenotype or/and summary statistics
- Input : genetics data and phenotype or/and summary statistics.
- Steps :
  - format and prepared files
  - build matrix of relatedness or/and genetic relationships matrix for GCTA, GEMMA
  - computed heritability using GEMMA and LDSC using summary statistics and GCTA, GEMMA and BOLT-LMM using genetics and phenotype.
- Test: using 4 phenotypes of lipid, genotype and summary statistics obtained using association testing result, we ran heritability and co-heritability using the cluster.
- Output : pipeline gave all intermediate file from each software but also a barplot with each heritability (see Figure 6)

An overview is shown in Figure 7 and the detailed computational cost is shown in Table 6.

| Process | Tot hours | % times | % cpu number used (Mean) | Max mem (MB) | NF processes |
|---|---|---|---|---|---|
| doGemmah2 | 4.10 | 4.26 | 372.19 | 4200.00 | 8 |
| doGemmah2_Stat | 0.00 | 0.01 | 67.39 | 43.00 | 8 |
| DoGemmah2Pval | 59.50 | 61.55 | 894.45 | 8300.00 | 4 |
| doGemmah2Pval_Stat | 0.00 | 0.00 | 63.38 | 43.00 | 4 |
| doGRLEM_GCTA_Stat_multi | 0.00 | 0.00 | 78.15 | 1.90 | 4 |
| doh2Bolt | 6.10 | 6.34 | 1523.67 | 5700.00 | 4 |
| doh2Bolt_Stat | 0.00 | 0.01 | 70.15 | 43.00 | 4 |
| doh2BoltiMulti | 17.00 | 17.61 | 1973.90 | 5700.00 | 1 |
| DoLDSC | 0.40 | 0.44 | 98.10 | 13900.00 | 1 |
| doLDSC_Stat | 0.00 | 0.00 | 29.50 | 43.00 | 1 |
| doMultiGRM | 0.50 | 0.55 | 779.73 | 5800.00 | 4 |
| GCTAComputeMultiGRM | 0.50 | 0.51 | 495.40 | 6500.00 | 1 |
| GCTAGRMByFile | 7.40 | 7.66 | 99.40 | 2700.00 | 4 |
| GCTAStrat | 0.00 | 0.00 | 79.00 | 158.00 | 1 |
| getBoltPhenosCovar | 0.00 | 0.01 | 47.60 | 65.30 | 1 |
| getGctaPhenosCovar | 0.00 | 0.04 | 44.80 | 66.60 | 4 |
| getGemmaRel | 0.80 | 0.82 | 673.60 | 5100.00 | 1 |
| MergeFile | 0.00 | 0.00 | 31.20 | | 1 |
| MergeH2 | 0.00 | 0.01 | 64.40 | 106.80 | 1 |
| select_rs_format | 0.20 | 0.17 | 54.40 | 609.70 | 1 |

Table 6: Statistics resume of heritability workflow running with cluster, using 4 phenotypes and 10,000 individuals and corresponding summary statistics. Process – Nextflow process name; Tot. hours – total hours used by NF process; % times – percentage of times used by process compared to other process ; % cpu number used (Mean) – mean % cpu number used by the process; Max mem (MB) is maximum of memory (RSS) used by any process; NF processes – number of Nextflow processes used for the steps.

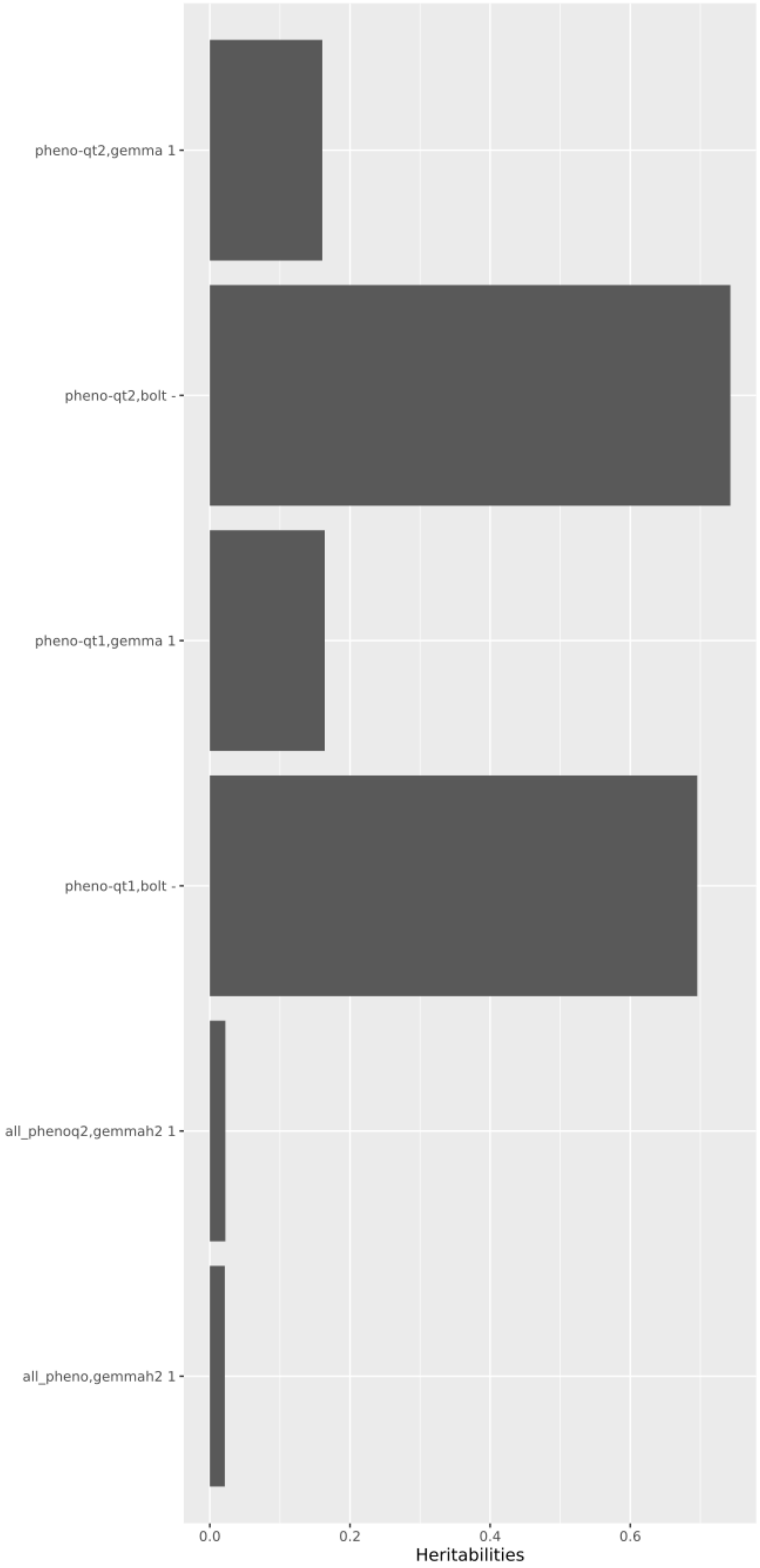

Figure 6: example of output of heritability workflow

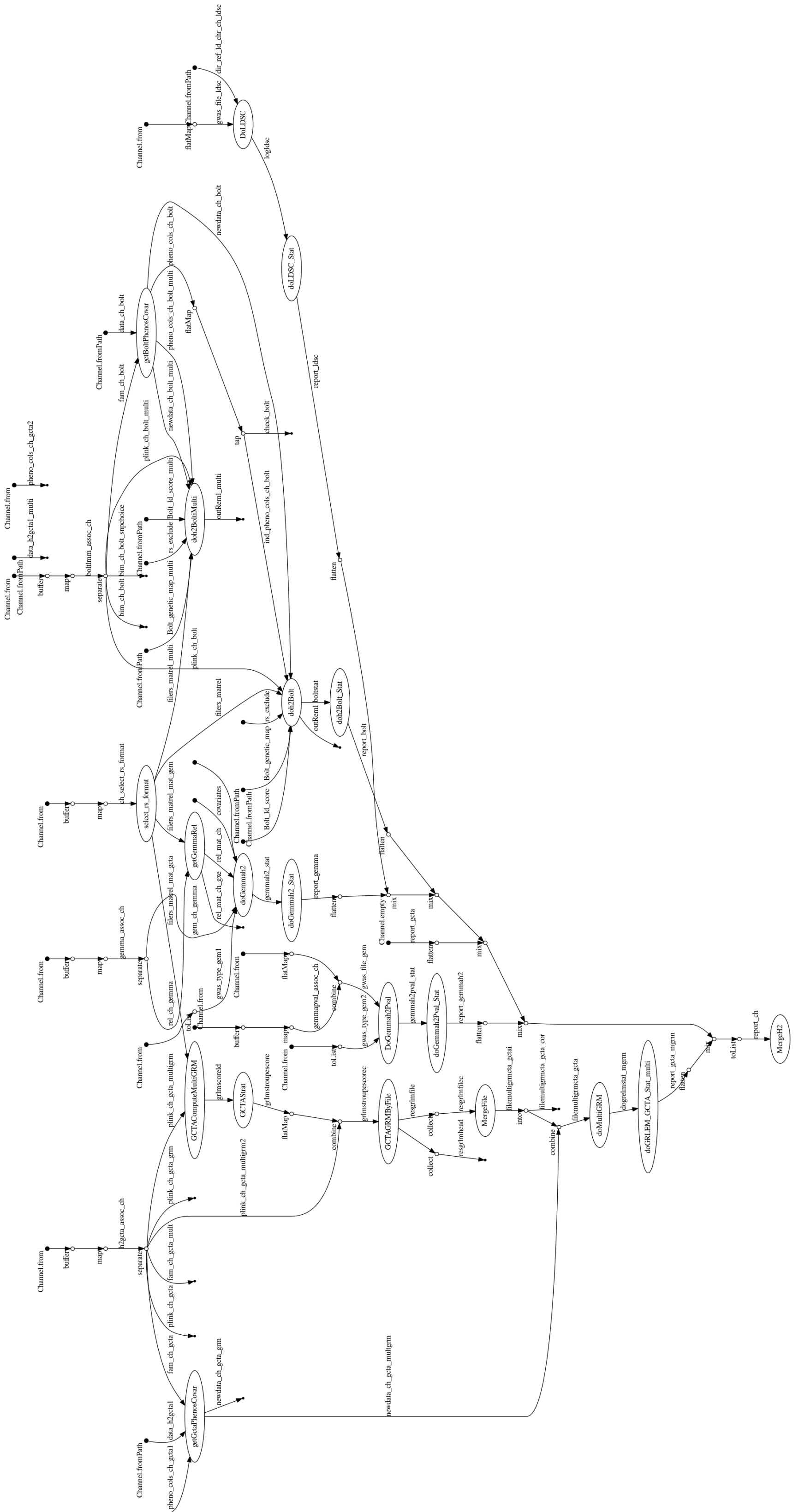

Figure 7: flowchart of heritability workflow

#### 3.5 Simulations workflow

Three workflows of simulation exist

##### 3.5.1 Description of each workflow

| Workflow | Input | genotype | Simulation process | Output | Post-simulation computation |
|---|---|---|---|---|---|
| `utils/build_example_data/main.nf` | • genotype in VCF file (or downloaded by ftp, by default 1000 Genomes, v37),<br>• effect database (or download by ftp GWAS catalog), phenotype of database,<br>• positions reference in BED format as array positions<br>• phenotype of effect database | Extract positions reference from genetics data, clean and format in plink file | extract positions and effect from the Effect database, extract corresponding genotype and simulated phenotypes with GCTA | • genetic data of population<br>• phenotypes quantitative or qualitative | Randomly switch the sex of some individuals to test `qc` pipeline |
| `utils/build_example_data/simul-assoc_gcta.nf` | • genotype in plink file<br>• effect database<br>• phenotype of effect database | None | as above | phenotypes quantitative or qualitative | None |
| `utils/build_example_data/simul-assoc_phenosim.nf` | genotype in plink file | None | random positions will be selected and phenotype simulated using *phenosim* | • phenotype simulated<br>• summary statistics of associations and statistics (FP and TP) | GEMMA and BOLT-LMM will run on genetics data and phenotype simulated. False Positive, True positive rate will be computed |

Table 7: Description : input - Input of different pipeline, genotype - if pipeline produce also a independent genotype complementary of phenotype simulation; simulation - how pipeline simulated phenotype; output - what output give pipeline; Post-simulation computation - Analyse or modification done after simulations

##### 3.5.2 Simulation using 1000Genome, GWAS catalog and GCTA

- Objectives : building phenotype using genetics data
- Input : by default, workflow uses (1) genetics data from 1000 Genomes Project (2) result of lead SNPs from GWAS catalog (3) list of phenotype choice in GWAS catalog to build phenotypes.
- Steps
  - Downloads GWAS catalog.

– Extracts and format GWAS catalog file with extraction of positions and effect using list of SNPs.
– Downloads genomics data of positions extracted from GWAS catalog and array.
– Extracts independent positions from position of GWAS catalog using "–clump" of PLINK and genetic data download.
– Uses Genomics data and $z$ values extracted from GWAS catalog using independent positions to build phenotype using GCTA.
– Output :
  * genotype in plink format of positions from array defined in input.
  * Quantitative and qualitative phenotype with position and genotype used to build phenotype corresponding and information relative to GWAS catalog and used to build phenotype.

- Test : build phenotypes using data of 1000 Genomes project, GWAS catalog and *diabetes* as phenotype.

See Table 8 for the detailed computational costs and Figure 8 for an overview of the different steps.

| Process | Tot hours | % times | % cpu number used (Mean) | Max mem (MB) | NF processes |
|---|---|---|---|---|---|
| addSexFile | 0.00 | 0.01 | 22.00 | 3.90 | 1 |
| cleanPLINKFile | 0.20 | 0.26 | 417.94 | 1.50 | 22 |
| cleanPLINKFile_GC | 0.40 | 0.56 | 307.69 | 1.50 | 22 |
| Dl1000G | 52.10 | 75.30 | 1.50 | 38.30 | 22 |
| Dl1000G_GC | 3.90 | 5.71 | 0.95 | 13.80 | 22 |
| format_sim_qualitatif | 0.00 | 0.03 | 90.30 | 20.90 | 1 |
| format_sim_quantitatif | 0.00 | 0.03 | 88.80 | 15.10 | 1 |
| format_simulated | 0.00 | 0.01 | 91.30 | 13.70 | 1 |
| getchr_gc | 0.00 | 0.03 | 39.30 | | 1 |
| GWASCatDl | 0.10 | 0.11 | 75.30 | 245.30 | 1 |
| mergePLINKFile | 0.00 | 0.01 | 135.00 | 71.20 | 1 |
| mergePLINKFile_GC | 0.00 | 0.01 | 307.40 | 12.70 | 1 |
| simulation_qualitatif | 0.00 | 0.01 | 89.10 | 1.50 | 1 |
| simulation_quantitatif | 0.00 | 0.01 | 95.00 | 5.60 | 1 |
| transfvcfInBed1000G | 2.10 | 3.11 | 422.98 | 9.20 | 22 |
| transfvcfInBed1000G_GC | 10.20 | 14.79 | 530.84 | 1.50 | 22 |

Table 8: Summary result for simulation individual using GWAS catalog for phenotype and 1000 genome project as genotype data. Process - it is Nextflow process name; Tot. hours – total hours used by NF process; % times – percentage of times used by process compared to other process ; % cpu number used (Mean) - mean % cpu number used by the process; Max mem (MB) is maximum of memory used by one process; NF processes - number of Nextflow process used for the steps

#### 3.6 Format data

##### 3.6.1 Convert PLINK format to VCF

- Objective : converts data to VCF for imputation

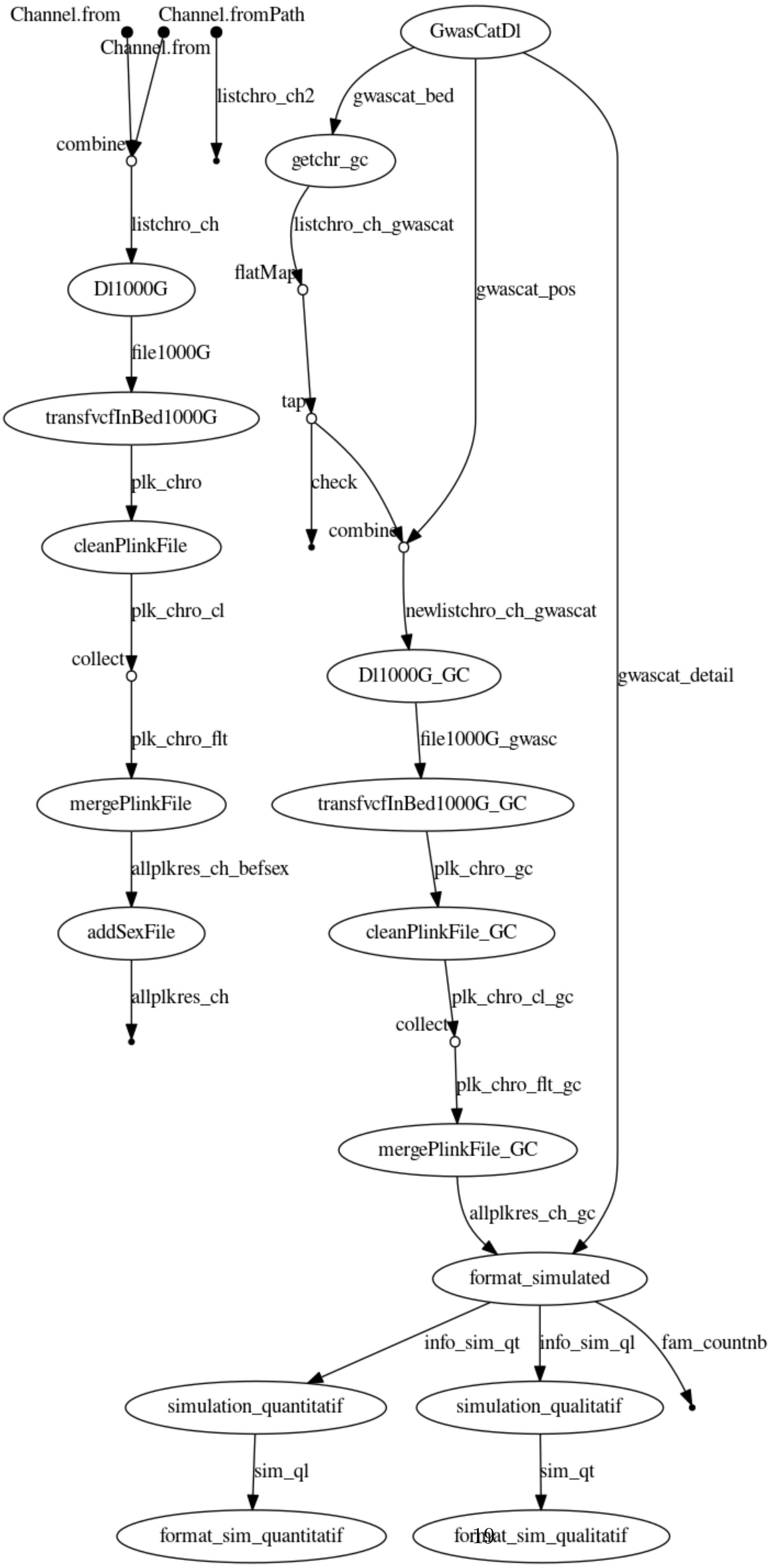

Figure 8: flowchart of simulation workflow

- Input PLINK file, reference genome of FASTA file and reference for positions, chromosome, and rs name.
- Steps
  - extract rsid and information of each positions from the reference file and reference sequence.
  - splits file by chromosome (optional)
  - converts PLINK to VCF.
  - cleans and rename position name.
  - fix allele using BCFtools
- Output : VCF file and VCF file by chromosome.
- Test : convert data after QC of 10,796 individuals using workflow in VCF format

See Figure 9 for an overview of the process and Table 9 for the detailed computational costs.

| Process | Tot hours | % times | % cpu number used (Mean) | Max mem (MB) | NF processes |
|---|---|---|---|---|---|
| checkfixref | 0.40 | 7.73 | 98.80 | 11.00 | 1 |
| checkVCF | 0.40 | 8.12 | 97.40 | 173.40 | 1 |
| convertInVcfChro | 3.00 | 62.03 | 155.99 | 1100.00 | 22 |
| convertrsname | 0.00 | 0.43 | 86.50 | 211.60 | 1 |
| CounChro | 0.10 | 1.29 | 3.50 | 2.60 | 1 |
| deletedmultianddel | 0.00 | 0.63 | 93.30 | 687.60 | 1 |
| extractpositionfasta | 0.00 | 0.19 | 50.90 | 1.50 | 1 |
| extractrsname | 0.70 | 13.46 | 98.60 | 505.60 | 1 |
| mergevcf | 0.20 | 3.86 | 319.00 | 16.00 | 1 |
| refallele | 0.10 | 2.26 | 13.80 | 165.60 | 1 |

Table 9: Summary of the result of the workflow to format PLINK to VCF prepare data for imputation. Process – Nextflow process name; Tot. hours – total hours used by NF process; % times – percentage of times used by process compared to other process; % cpu number used (Mean) – mean % cpu number used by the process; Max mem (MB) is maximum of memory used by one process (resident set size); NF processes – number of Nextflow processes used for the steps

###### 3.6.2 Convert VCF to PLINK or other format

- Objective : convert output from imputation in PLINK or other format to run association testing.
- Input : list of VCF, genetic map.
- Steps
  - Computed various statistics as frequency and imputation score
  - Filter VCF by score and frequency
  - Transform file to PLINK.
  - check for rs duplicate and correct.

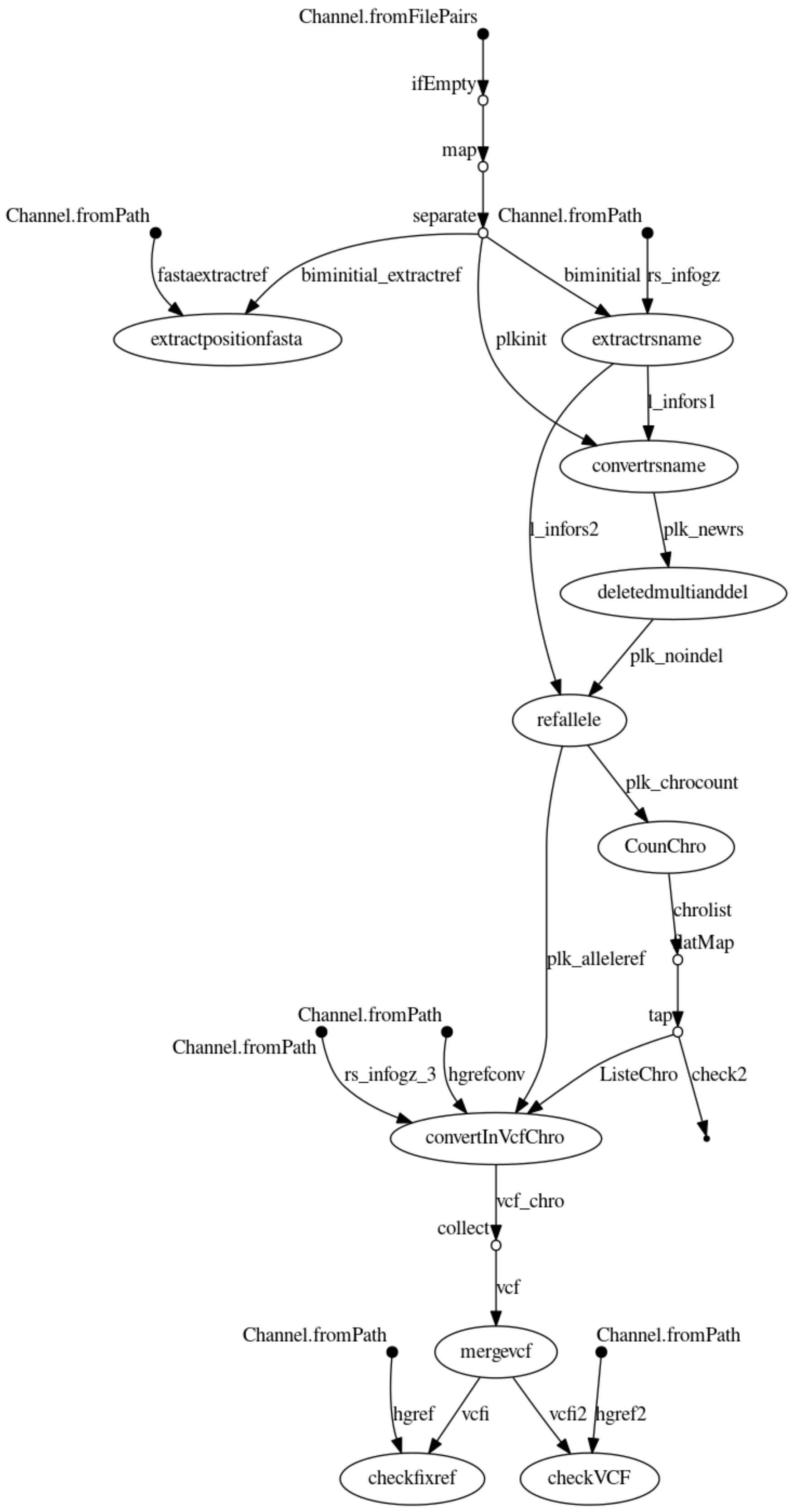

Figure 9: Flowchart of workflow to convert PLINK format to VCF to prepare data for imputation

– merge all files PLINK by chromosome.

- Output: PLINK file and report presenting distribution of quality and frequencies 10 and plink files converted.
- Test : used output of imputed data (≈30 millions SNPs and 12,000 individuals)

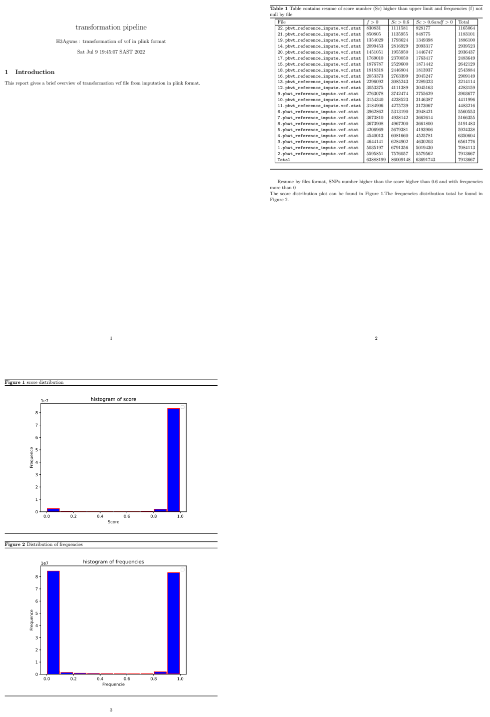

transformation pipeline

H3Agwas : transformation of vcf in plink format

Sat Jul 9 19:45:07 SAST 2022

1 Introduction

This report gives a brief overview of transformation vcf file from imputation in plink format.

1

**Table 1** Table contains resume of score number (Sc) higher than upper limit and frequencies (f) not null by file

| File | $f > 0$ | $Sc > 0.6$ | $Sc > 0.6 and f > 0$ | Total |
|---|---|---|---|---|
| 22.pbwt_reference_impute.vcf.stat | 830831 | 1111581 | 828177 | 1165064 |
| 21.pbwt_reference_impute.vcf.stat | 850805 | 1135955 | 848775 | 1183101 |
| 19.pbwt_reference_impute.vcf.stat | 1354029 | 1793624 | 1349398 | 1886100 |
| 14.pbwt_reference_impute.vcf.stat | 2099453 | 2816929 | 2093317 | 2939523 |
| 20.pbwt_reference_impute.vcf.stat | 1451051 | 1955950 | 1446747 | 2036437 |
| 17.pbwt_reference_impute.vcf.stat | 1769010 | 2370050 | 1763417 | 2483649 |
| 15.pbwt_reference_impute.vcf.stat | 1876787 | 2529600 | 1871442 | 2642129 |
| 18.pbwt_reference_impute.vcf.stat | 1818318 | 2446804 | 1813937 | 2543884 |
| 16.pbwt_reference_impute.vcf.stat | 2053373 | 2763399 | 2045247 | 2909149 |
| 13.pbwt_reference_impute.vcf.stat | 2296092 | 3085243 | 2289323 | 3214114 |
| 12.pbwt_reference_impute.vcf.stat | 3053375 | 4111389 | 3045163 | 4283159 |
| 9.pbwt_reference_impute.vcf.stat | 2763078 | 3742474 | 2755629 | 3903677 |
| 10.pbwt_reference_impute.vcf.stat | 3154340 | 4238523 | 3146387 | 4411996 |
| 11.pbwt_reference_impute.vcf.stat | 3184906 | 4275739 | 3173067 | 4483216 |
| 6.pbwt_reference_impute.vcf.stat | 3962862 | 5313190 | 3948421 | 5560553 |
| 7.pbwt_reference_impute.vcf.stat | 3673810 | 4938142 | 3662614 | 5166355 |
| 8.pbwt_reference_impute.vcf.stat | 3673908 | 4967200 | 3661800 | 5191483 |
| 5.pbwt_reference_impute.vcf.stat | 4206969 | 5679381 | 4193906 | 5924338 |
| 4.pbwt_reference_impute.vcf.stat | 4540013 | 6081660 | 4525781 | 6350604 |
| 3.pbwt_reference_impute.vcf.stat | 4644141 | 6284902 | 4630203 | 6561776 |
| 1.pbwt_reference_impute.vcf.stat | 5035197 | 6791356 | 5019430 | 7084113 |
| 2.pbwt_reference_impute.vcf.stat | 5595851 | 7576057 | 5579562 | 7913667 |
| Total | 63888199 | 86009148 | 63691743 | 7913667 |

Resume by files format, SNPs number higher than the score higher than 0.6 and with frequencies more than 0
The score distribution plot can be found in Figure 1.The frequencies distribution total be found in Figure 2.

2

**Figure 1** score distribution

**Figure 2** Distribution of frequencies

3

Figure 10: Output of pipeline format vcf in plink, with distribution of quality and frequency (figures with general distribution and table distribution by frequency)

An overview is shown in Figure 11 and the detailed computational costs are shown in Table 10.

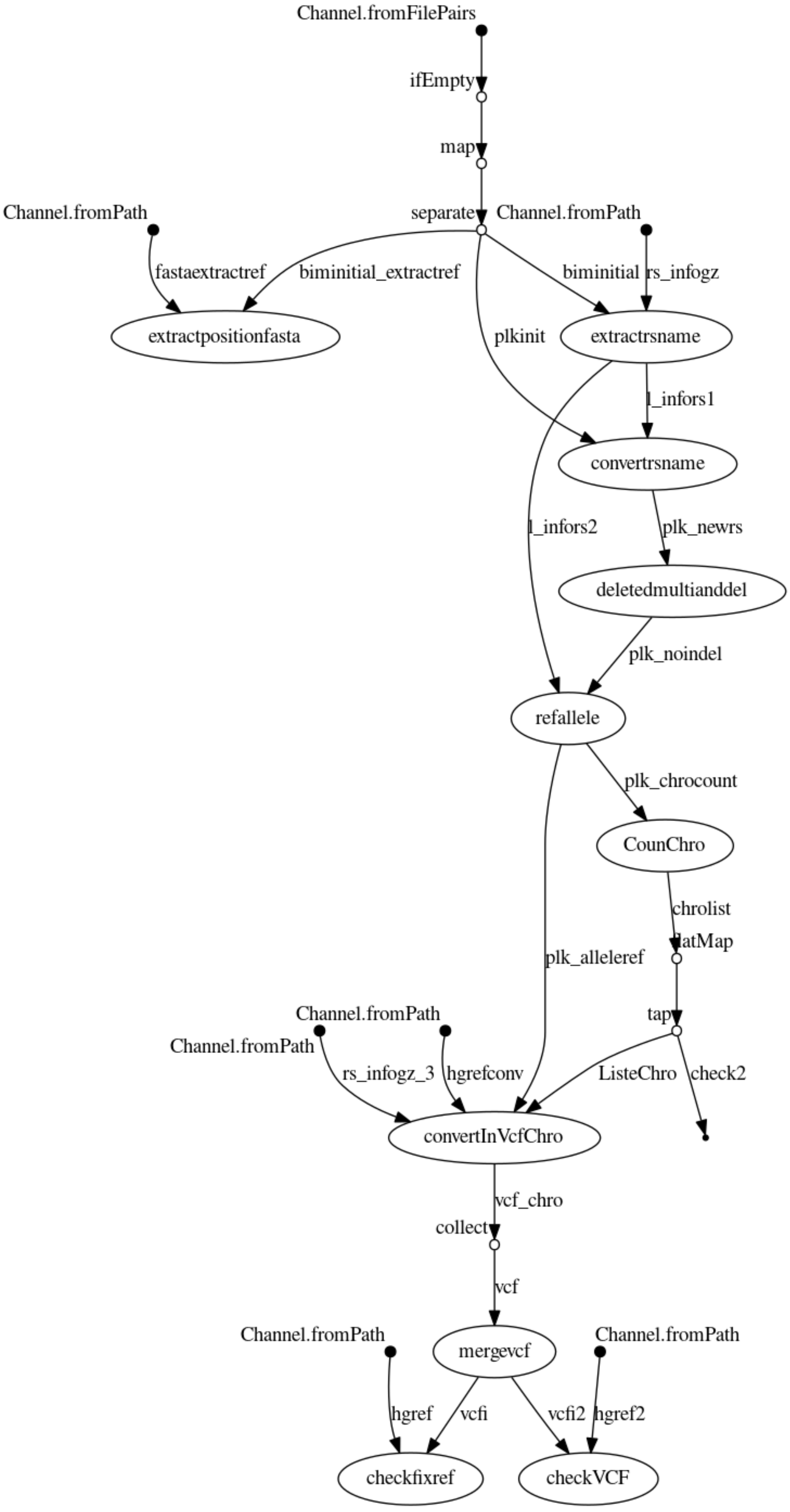

Figure 11: flowchart of workflow to convert VCF file from imputation into PLINK format

| Process | Tot hours | % times | % cpu number used (Mean) | Max mem (MB) | NF processes |
|---|---|---|---|---|---|
| TransformRsDup | 7.20 | 4.71 | 10.97 | 224.70 | 22 |
| AddedCM | 2.80 | 1.83 | 26.99 | 187.50 | 22 |
| computedstat | 13.20 | 8.65 | 99.33 | 8.60 | 22 |
| dostat | 0.10 | 0.08 | 96.80 | 6400.00 | 1 |
| formatvcfscore | 128.00 | 84.08 | 108.37 | 196.10 | 22 |
| GetRsDup | 0.00 | 0.02 | 89.00 | 2900.00 | 1 |
| MergePLINK | 1.00 | 0.64 | 83.10 | 62700.00 | 1 |

Table 10: Summary of result of workflow to format VCF after imputation in PLINK format using data after imputation obtained in QC. Process – Nextflow process name; Tot. hours – total hours used by NF process; % times – percentage of times used by process compared to other process ; % cpu number used (Mean) – mean % cpu number used by the process; Max mem (MB) is maximum of memory used by one process; NF processes – number of Nextflow processes used for the steps.

##### 3.6.3 Multi-trait analyse using MTAG

- Objective : Analyse multi trait using summary statistics
- Input : result of summary statistics from various phenotype
- Steps
  - format each files to prepared input for MTAG software
  - Run MTAG with all summary statistics
  - Run MTAG seletected 2 by 2 each summary statistics.
- Output : result of mtag sofware (summary statistics), and report in PDF as association pipeline
- Test : used output summary statistics (14 millions SNPs) for 4 SNPs

An overview is shown in Figure 12 and the detailed computational costs in Table 11.

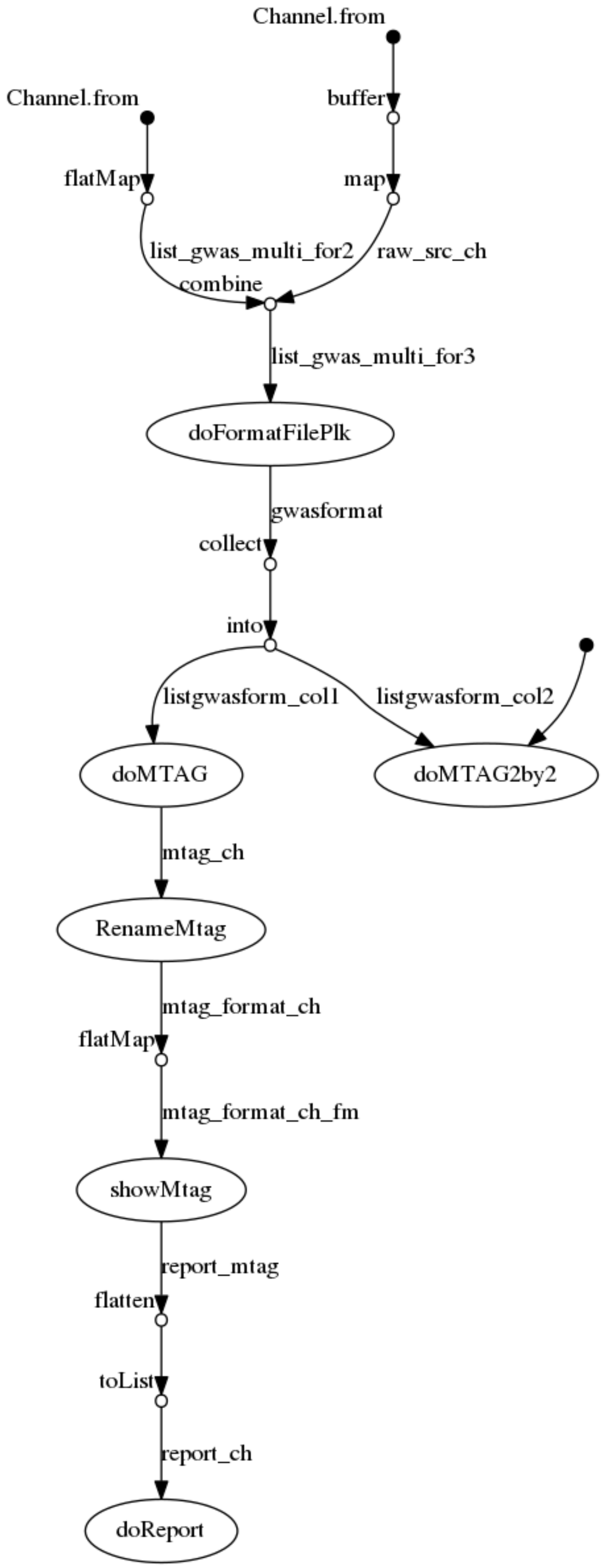

Figure 12: flowchart of workflow to convert VCF file from imputation in PLINK format

| Process | Tot hours | % times | % cpu number used (Mean) | Max mem (MB) | NF processes |
|---|---|---|---|---|---|
| doFormatFilePlk | 0.70 | 13.13 | 71.35 | 6400.00 | 4 |
| doMTAG | 1.20 | 22.82 | 98.70 | 34600.00 | 1 |
| doMTAG2by2 | 3.00 | 59.39 | 100.35 | 18600.00 | 6 |
| doReport | 0.00 | 0.02 | 45.10 | 20.50 | 1 |
| RenameMtag | 0.00 | 0.88 | 17.70 | 9.30 | 1 |
| showMtag | 0.20 | 3.76 | 91.75 | 6900.00 | 4 |

Table 11: Summary of result of workflow doing a multi trait analysis using MTAG: Process – Nextflow process name; Tot. hours – total hours used by NF process; % times – percentage of times used by process compared to other process; % cpu number used (Mean) – mean % cpu number used by the process; Max mem (MB) is maximum of memory used by one process; NF processes – number of Nextflow processes used for the steps.

#### 4 Detail of experimentation in the main paper

These tables give detail of the experimentation described in the main paper.

| Process | Tot hours | % times | % cpu number used (Mean) | Max mem (MB) | NF processes |
|---|---|---|---|---|---|
| computePCA | 2.3 | 0.5 | 374 | 1100 | 1 |
| drawPCA | 0.0 | 0.0 | 114 | 89 | 1 |
| extractPheno | 0.0 | 0.0 | 81 | 63 | 1 |
| bgen_formatsample | 0.0 | 0.0 | 92 | 93 | 1 |
| indexbgen_list | 3.4 | 0.8 | 41 | 7 | 22 |
| getBoltPhenosCovar | 0.0 | 0.0 | 79 | 68 | 1 |
| select_rs_format | 0.0 | 0.0 | 58 | 635 | 1 |
| FastGWADoGRM | 11.5 | 2.7 | 227 | 2100 | 100 |
| MergFastGWADoGRM | 0.0 | 0.0 | 17 | 3 | 1 |
| computeTest | 8.1 | 1.9 | 100 | 824 | 4 |
| format_genetic_ldscore | 0.4 | 0.1 | 43 | 2200 | 1 |
| doBoltmm | 5.7 | 1.3 | 351 | 3600 | 4 |
| getListeChroGem | 0.0 | 0.0 | 120 | 5 | 1 |
| getGemmaRel | 0.6 | 0.1 | 854 | 5100 | 1 |
| doGemmaChro | 312.1 | 72.7 | 945 | 5400 | 88 |
| doMergeGemma | 0.1 | 0.0 | 24 | 4 | 4 |
| FastGWARun | 41.9 | 9.8 | 972 | 2200 | 4 |
| getListeChro_saige | 0.0 | 0.0 | 81 | 5 | 1 |
| getchrobgen | 0.0 | 0.0 | 86 | 23 | 22 |
| getSaigePheno | 0.0 | 0.0 | 103 | 66 | 1 |
| checkidd_saige | 0.0 | 0.0 | 43 | 136 | 1 |
| subplink_heritability_saige | 0.0 | 0.0 | 74 | 927 | 1 |
| saige_computed_variance | 1.5 | 0.4 | 878 | 806 | 4 |
| doSaigeListBgen | 13.4 | 3.1 | 99 | 644 | 88 |
| doMergeSaige | 0.0 | 0.0 | 40 | 4 | 4 |
| regenie_step1 | 4.6 | 1.1 | 176 | 5900 | 4 |
| regenie_step2 | 22.6 | 5.3 | 925 | 373 | 88 |
| merge_regenie | 0.1 | 0.0 | 24 | 4 | 4 |
| format_regeniesumstat | 0.1 | 0.0 | 78 | 10 | 4 |
| ShowManhattan | 0.7 | 0.2 | 97 | 7900 | 20 |
| drawPlinkResults | 0.1 | 0.0 | 83 | 4300 | 4 |
| showPhenoDistrib | 0.0 | 0.0 | 85 | 127 | 1 |
| doReport | 0.0 | 0.0 | 34 | 25 | 1 |

Table 12: Cost of associationon cluster, using 4 phenotypes and 10,700 individuals. The elapsed time for the entire workflow was 12h 36min, with a high degree of parallelisation. *Process* is the Nextflow process name; *Tot hours* – total CPU hours used by instances of this NF process; *% times* – % of time used by process compared to other process; *% cpu number used (Mean)* – mean % cpu number used by instances of the process — a measure of achievable parallelism for instances of that process; *Max mem (MB)* is the maximum resident set size used by one process; *NF processes* – number of Nextflow process used for the steps, and a measure of parallelism at very coarse level.

| Process | Tot hours | % times | % cpu number used (Mean) | Max mem (MB) | NF processes |
|---|---|---|---|---|---|
| GetRsFile | 1.1 | 1.2 | 99.5 | 9 | 1 |
| ChangeFormatFile | 29.9 | 30.7 | 65.2 | 4100 | 3 |
| doGWAMA | 18.0 | 18.5 | 67.4 | 10400 | 1 |
| doMetal | 2.9 | 3.0 | 99.6 | 3500 | 1 |
| doMetaSoft | 12.3 | 12.7 | 103.1 | 7700 | 1 |
| doMRMEGA | 15.8 | 16.2 | 68.1 | 8100 | 1 |
| doPlinkMeta | 8.9 | 9.2 | 99.8 | 2000 | 1 |
| showGWAMA | 1.4 | 1.5 | 100.2 | 5800 | 1 |
| showMetal | 1.8 | 1.8 | 97.7 | 3800 | 1 |
| showMetasoft | 1.9 | 1.9 | 100.1 | 7500 | 1 |
| showMRMEGA | 1.6 | 1.6 | 100.2 | 6200 | 1 |
| showPlink | 1.2 | 1.3 | 100.0 | 4800 | 1 |
| doReport | 0.6 | 0.6 | 46.1 | 24 | 1 |

Table 13: Cost of running meta-analysis workflow using Wits cluster, using 3 summary statistics and 14 millions of positions by summary statistics. Column labels as in Table 12
